# Powerful and accurate detection of temporal gene expression patterns from multi-sample multi-stage single cell transcriptomics data with TDEseq

**DOI:** 10.1101/2023.09.12.557282

**Authors:** Yue Fan, Lei Li, Shiquan Sun

## Abstract

We present a non-parametric statistical method called TDEseq that takes full advantage of smoothing splines basis functions to account for the dependence of multiple time points, and uses hierarchical structure linear additive mixed models to model the correlated cells within an individual. As a result, TDEseq demonstrates powerful performance in identifying four potential temporal expression patterns within a specific cell type. Extensive simulation studies and the analysis of four published scRNA-seq datasets show that TDEseq can produce well-calibrated p-values and up to 20% power gain over the existing methods for detecting temporal gene expression patterns.

## INTRODUCTION

The advances in single-cell RNA sequencing (scRNA-seq) technologies make it possible to record the temporal dynamics of gene expression over multiple time points or stages either in the same cell population [1, 2] or even in an individual cell without destruction [3]. Unlike the single time point (e.g., snapshot) profiling of transcriptome that allocates cells on pseudotime or lineages using purely computational strategies [4-6], in particular, the time-course scRNA-seq profiling of whole transcriptome with respect to real, physical time, is capable of providing additional insights into dynamic biological processes [2, 7]. For example, how the cells naturally differentiate into other types or states during the development processes and how the cellular response to specific drug treatments [8], viral infections [9], etc. Therefore, accurately characterizing the temporal dynamics of gene expression over time points is crucial for developmental biology [10, 11], tumor biology [12-14], and biogerontology [15-17], which allows us to decipher the dynamic cellular heterogeneity during cell differentiation [18]; identifying cancer driver genes during the status transformation [14]; and investigating the mechanisms of cell senescence during aging [15]. Although the time-course scRNA-seq studies are initially designed for different purposes, they essentially require the same data analysis tools for detecting the temporal dynamics of gene expression [19]. As we know, the statistical modeling for this type of scRNA-seq data to identify temporal gene expression patterns meets significant challenges, i.e., modeling unwanted variables, accounting for temporal dependencies, and even characterizing non-stationary cell populations of scRNA-seq data. However, existing methods are unable to fully consider these limitations, resulting it remains an urgent need to develop effective tools for detecting temporal dynamics of gene expression in time-course scRNA-seq studies.

Particularly, time-course scRNA-seq data commonly share a fundamental temporal dynamics nature, i.e., the gene expression levels measured at each time point may be influenced by previous time points. Accounting for these temporal dependencies requires specialized statistical and computational tools [20], and failure to do so can lead to inaccurate gene detections [21, 22]. As a result, current temporal gene detection methods for time-course scRNA-seq data can be divided into two categories: the methods that treat time points independently and methods that model the temporal dependencies explicitly. Specifically, the methods that utilize the former approach mostly treated time as a categorical variable, performing the differential expression analysis with pair-wise comparison tools, such as a two-sided *Wilcoxon* rank-sum test [23, 24], etc. However, neglecting the temporal dependencies among multiple time points will reduce the statistical power and may lead to false-positive results [22]. On the other hand, the methods that utilize the latter approach are commonly used for addressing the time-course bulk RNA-seq data, such as ImpulseDE2 [25], DESeq2 [26], and edgeR [27]. However, the scRNA-seq data is often sparse along with technical and biological variability, making it difficult to accurately identify true biological gene expression changes over multiple time points [28, 29].

Furthermore, time-course scRNA-seq data are often collected from multi-sample multi-stage designs. As a result, there may be unwanted variables that arise due to technical variability, batch effects, or the genetic background of individuals [30]. These variables can obscure the identification of temporal expression changes that are of interest, making it challenging to detect temporal expression genes accurately. Alternatively, the trajectory-based differential expression analysis methods, such as Monocle2 [5], tradeSeq [31], and PseudotimeDE [32] could detect the temporal dynamics of gene expressions along with pseudotime or a continuous trajectory of cellular states. However, since the gene expression profiles of cells from the same sample/individual are known to be dependent, these methods may not adequately account for technical or biological variability that may present in multi-sample multi-stage designs. In addition, these methods may not fully capture the underlying biological process at specific tipping points or intervals, which could be particularly relevant in understanding the mechanisms of cell fate or differentiation [33, 34], and tumor progression [14].

Here, to properly address the above challenges, we develop an efficient and flexible non-parametric method for detecting temporal expression patterns over multiple time points. We refer to our method as *TDEseq*, Temporal Differentially Expressed genes of time-course scRNA-seq data. Specifically, TDEseq primarily builds upon a linear additive mixed model (LAMM) framework, with a random effect term to account for correlated cells in time-course scRNA-seq studies. In this model, we typically introduce the quadratic *I*-splines [35] and cubic *C*-splines [36] as basis functions, which facilitate the detection of four potential temporal gene expression patterns, i.e., growth, recession, peak, or trough. As a result, with extensive simulation studies, we find TDEseq can properly control for type I error rate at the transcriptome-wide level, and display powerful performance in detecting temporal expression genes under the power simulations. Finally, we apply the TDEseq to one scRNA-seq data set which was generated by Well-TEMP-seq [23], one scRNA-seq data set generated by Smart-seq2 [37], and two scRNA-seq data sets generated by 10X Genomics, to benchmark TDEseq against current state-of-the-art methods, regarding human colorectal cancer development [23], mouse hepatocyte differentiation [38], human metastatic lung adenocarcinoma [14] and human COVID-19 progression [9]. These results highlight that TDEseq is an appropriate tool for detecting temporal gene expression patterns over multiple time points, which leads to an improved understanding of developmental biology, tumor biology, and biogerontology.

## RESULTS

### Overview of TDEseq

#### Statistical modeling

TDEseq is a temporal gene expression analysis approach that is primarily built upon the linear additive mixed models (LAMM) [39] framework to characterize the temporal gene expression changes for time-course (or longitudinal) scRNA-seq data sets (Materials and Methods; Supplementary Text). Typically, we aim to detect one of four possible temporal gene expression patterns (i.e., growth, recession, peak, or trough) over multiple time points using both *I*-splines [35] and *C*-splines [36] basis functions (Fig. 1A), and examine one gene at a time. Briefly, In LAMM, we assume the log-normalized gene expression level of raw counts (Materials and Methods), i.e., *y_gji_*(*t*), for gene *g*, individual *j* and cell *i*, at time point *t*, is,

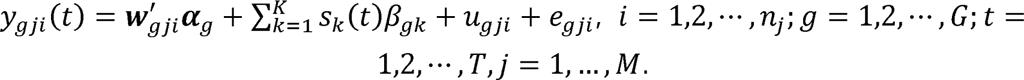

**Fig. 1.**
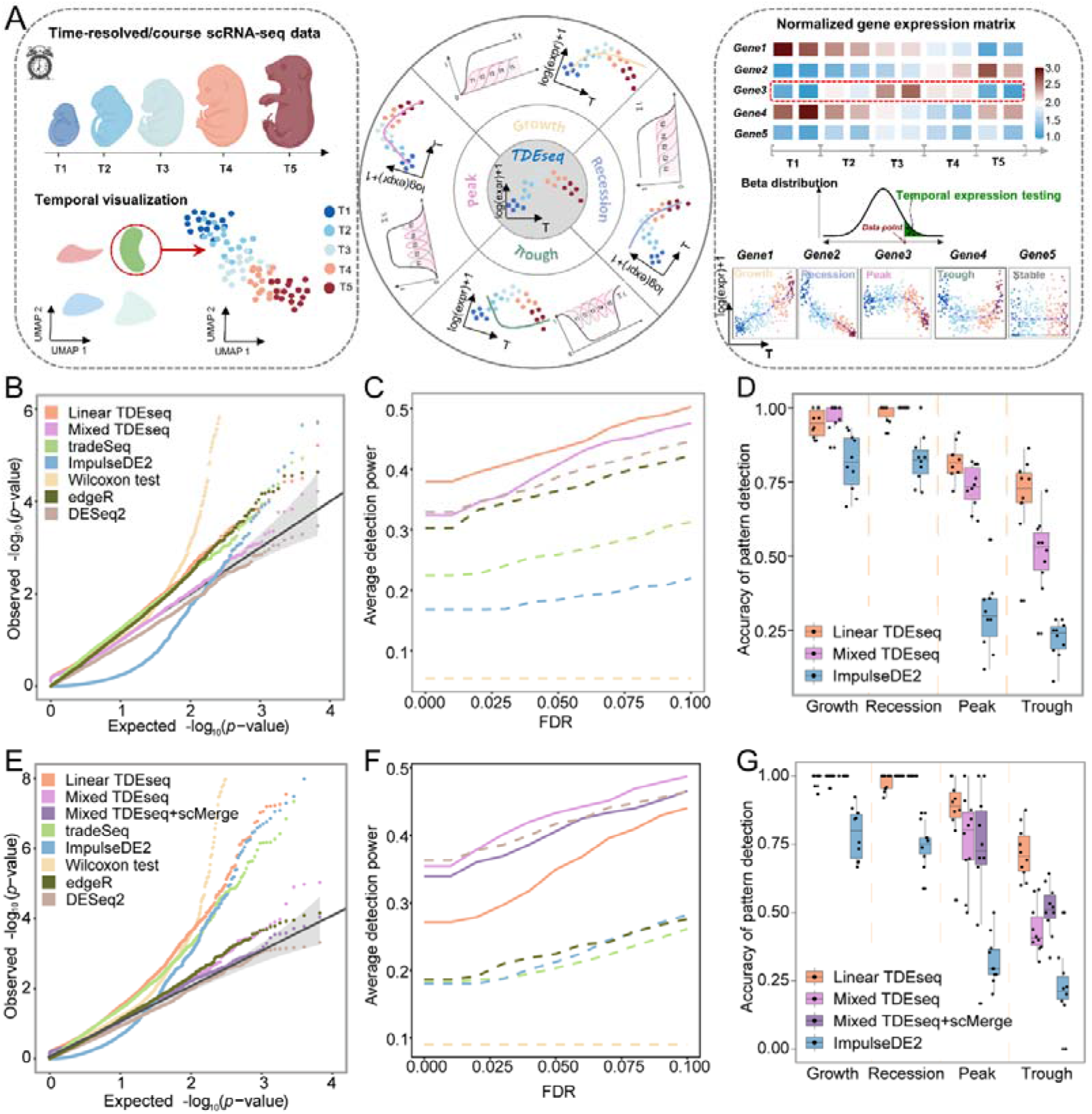
Schematic overview of TDEseq and the methods comparison in simulations. (**A**) TDEseq is designed to perform temporal expression gene analysis on time-course scRNA-seq data. With a given gene, TDEseq determines one of four temporal expression patterns, i.e., growth, recession, peak, and trough. TDEseq combines the four *p*-values using the Cauchy combination rule as a final *p*-value, facilitating the detection of one of four temporal gene expression patterns. (**B**) The quantile-quantile (QQ) plot shows the type I error control under the baseline parameter settings. The well-calibrated *p*-values will be expected laid on the diagonal line. The *p-*values generated from Mixed TDEseq (plum) and DESeq2 (brown) are reasonably well-calibrated, while Linear TDEseq (orange), tradeSeq (green), ImpulseDE2 (blue), Wilcoxon test (yellow) and edgeR (dark green) produced the *p*-values that are not well-calibrated. (**C**) The average power of 10 simulations for temporal expression gene detection across a range of FDR cutoffs under the baseline parameter settings. Both versions of TDEseq exhibit high detection power of temporal expression genes, followed by DESeq2, edgeR, tradeSeq, and ImpulseDE2. Wilcoxon test does not fare well, presumably due to bias towards highly expressed genes. The TDEseq methods were highlighted using solid lines, while other methods were represented by dashed lines in the plots. (**D**) The comparison of Linear TDEseq, Mixed TDEseq, and ImpuseDE2 in terms of the accuracy of temporal expression pattern detection under the baseline parameter settings, at an FDR of 5%. The temporal expression genes detected by TDEseq demonstrated a higher accuracy than those detected by ImpluseDE2. (**E**) The quantile-quantile (QQ) plot shows the type I error control under the large batch effect parameter settings. The *p-*values generated from Mixed TDEseq coupled with scMerge (purple) and DESeq2 (brown) are reasonably well-calibrated, while Linear TDEseq (orange), Mixed TDEseq (plum), tradeSeq (green), ImpulseDE2 (blue), Wilcoxon test (yellow) and edgeR (dark green) generated the inflated *p*-values. (**F**) The average power of 10 simulations replicates the comparison of temporal expression gene detection across a range of FDR cutoffs under the large batch effect parameter settings. (**G**) The comparison of Linear TDEseq, Mixed TDEseq, and Mixed TDEseq coupled with scMerge and ImpuseDE2 in terms of the accuracy of temporal expression pattern detection under the large batch effect parameter settings, at an FDR of 5%. Since DESeq2, edgeR, tradeSeq, and Wilcoxon tests were not originally designed for pattern-specific detection we excluded them in the comparison. FDR denotes the false discovery rate.

where ***w**_gji_* is the cell-level or time-level covariate (e.g., cell size, or sequencing read depth), *α_g_* is its corresponding coefficient; ***u**_g_* is a random vector to account for the variations from heterogeneous samples, i.e.,

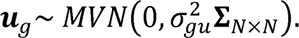

where **Σ**_*N×N*_ is a block diagonal matrix with a total of block matrices, in which all elements of **Σ**_*n_j_×n_j_*_ are ones; *n_j_* is the number of cells for the individual or replicate *j* and 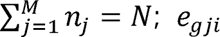 is a random effect, which is an independent and identically distributed variable that follows a normal distribution with mean zero and variance 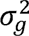 to account for independent noise, i.e.,

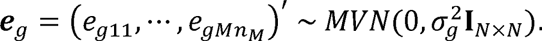

Particularly, the variable *s_k_*(*t*) is a smoothing spline basis function, which involves either *I*-splines or *C*-splines to model monotone patterns (i.e., growth and recession) and quadratic patterns (i.e., peak and trough), respectively [40] (Fig. 1A). The *I*-splines are defined as 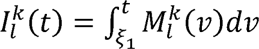 [35]; while *C*-splines are defined as 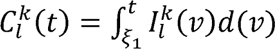, [36] based on *I*-splines, and *β_gk_* is its corresponding coefficient that is restricted to *β_gk_* ≥ 0, and *l* (*l* = 1,2, ..., *L*) is the number of grid points. We set *L* to be a total number of grid points in scRNA-seq data, which is equal to the number of time points; *k* denotes the order of the spline function; *MVN* denotes the multivariate normal distribution.

**Hypothesis testing**: in the LAMM model mentioned above, we are interested in examining whether a gene shows one of four temporal expression patterns, i.e., growth, recession, peak, or trough (Fig. 1A). Testing whether a gene expression hypothesis ***H**_0_:**β**_g_* = **0**. Parameter estimates and hypothesis testing in LAMM are displays temporal gene expression patterns can be translated into testing the null notoriously difficult, as the LAMM likelihood involves *M*-splines [35] (non-linear) subject to nonnegative constraints that cannot be solved analytically. To make the LAMM model scalable estimation and inference, we developed an approximate inference algorithm based on a cone programming projection algorithm [41, 42]. With parameter estimates, we computed a *p*-value for each of the four patterns using the test statistics [43], which follow a mixture of beta distributions [44]. Afterward, we combined these four *p*-values through the Cauchy combination rule [45]. The Cauchy combination rule allows us to combine multiple potentially correlated *p*-values into a single *p*-value to determine whether a gene exhibits the temporal expression pattern or not (Materials and Methods; Additional file 1: Supplementary Text).

We refer to the above method as the mixed version of TDEseq (we denoted as *Mixed TDEseq*). Besides the mixed version, we have also developed a linear version of TDEseq (to distinguish Mixed TDEseq, we denoted this as *Linear TDEseq*) for modeling the small or no sample heterogeneity inherited in time-course scRNA-seq data (Additional file 1: Supplementary Text). Both versions of TDEseq were implemented in the same R package with multiple threads computing capability. The software TDEseq, together with the reproducibility analysis code is freely available at https://sqsun.github.io/software.html.

### TDEseq generates well-calibrated *p*-values and exhibits powerful gene detection of temporal expression changes in simulations

To benchmark the robustness and performance of TDEseq, we simulated extensive scRNA-seq data sets using the *Splatter* R package [46], and compared two versions of TDEseq with other five existing approaches but not specific designs for time-course scRNA-seq data analysis, which are the two-sided Wilcoxon rank-sum test (Wilcoxon test), tradeSeq [47], ImpulseDE2 [25], edgeR [48, 49], and DESeq2 [26] (Materials and Methods). The simulations were typically designed to assess the ability of TDEseq in terms of type I error control and temporal gene detection power with varying various parameter settings, including the number of time points (i.e., 4, 5, or 6), the number of cells for each sample in each time point (i.e., 50, 100, or 200; three replicates/samples for each time point), the expected UMI counts for each cell of scRNA-seq data (i.e., 7.0 as low, 9.7 as medium, and 13.8 as high), the effect size of temporal expression changes (i.e., 0.1 as low, 0.4 as medium, and 0.7 as high), and the sample-level unwanted technical variations (i.e., batch effects; 0 as no batch effects, 0.04 as medium, and 0.12 as high).

To do so, we considered a baseline simulation scenario: the number of time points as 5; the number of cells in each sample as 300; the expected UMI counts for each cell as 9.7; the batch effect size as 0.04; the time point-specific effect size as 0.4; all cells were measured by 10,000 genes, in which 1,000 genes were randomly assigned one of four possible temporal patterns (i.e., growth, recession, peak, and trough; Additional file 2: Fig. S1A-S1D) in power simulations. With the baseline parameter settings, we varied one parameter at a time to examine whether the gene was temporally expressed over multiple time points. Notably, the expected UMI counts under baseline settings were estimated from the lung adenocarcinoma progression scRNA-seq data [14] (Materials and Methods).

With the baseline parameter setup, we found that only Mixed TDEseq and DESeq2 generated the well-calibrated *p*-values under the null simulations, whereas all other methods produced the inflated or conserved *p*-values (Fig. 1B). Besides, for the power simulations, Linear TDEseq and Mixed TDEseq can produce a more powerful temporal expression pattern detection rate across a range of FDR cutoffs (Fig. 1C). Specifically, with a false discovery rate (FDR) of 5%, the power detection rate of both Linear and Mixed TDEseq was 43.3% and 40.8%, respectively, followed by DESeq2 was 38.7%, edgeR was 36.4%, tradeSeq was 25.9% and ImpulseDE2 was 18.4%. Furthermore, we also examined the accuracy of pattern detection, finding both Linear and Mixed versions of TDEseq outperformed ImpluseDE2 (the sole method capable of identifying pattern-specific genes). Specifically, with an FDR of 5%, the averaged accuracy of pattern examination (with 10-time repeats) for Mixed TDEseq achieved 99.0% for growth, 100% for recession, 80.4% for peak, and 43.6% for trough. In contrast, ImpulseDE2 achieved 73.1% for growth, 83.5% for recession, 39.3% for peak, and 21.3% for trough (Fig. 1D).

In addition, we systematically examined the performance of the type I error control rate under other parameter settings. Our findings indicate that Mixed TDEseq consistently produces well-calibrated *p*-values (Additional file 2: Fig. S2A, S2B, S3A, S4A, and S4B) except when dealing with high UMI counts (Additional file 2: Fig. S3B). These observations are presumably due to the presence of sample-level variations or batch effects associated with high UMI counts. On the other hand, in terms of temporal expression gene detection and averaged accuracy of pattern examination, either Mixed or Linear TDEseq displayed more powerful performance across a range of parameter settings regardless of the number of time points (Additional file 2: Fig. S2C and S2D), the low expected UMI counts for each cell (Additional file 2: Fig. S3C), the large number of cells per sample (Additional file 2: Fig. S4D) and the small effect size setups (Additional file 2: Fig. S5), as well as the accuracy of pattern examination (Additional file 2: Fig. S6). Meanwhile, we found both pseudobulk-based methods, i.e., either edgeR or DESeq2, performed well with high UMI counts (Additional file 2: Fig. S2D) and small number of cells per sample (Additional file 2: Fig. S3C). These observations were probably consistent with the previous studies that DESeq2/edgeR performed well on log-normal distributed small sample size RNA-seq data [50, 51]. Taken together, we summarized the findings on detection power at a 5% FDR across diverse parameter settings. The results demonstrated that the power of temporal gene detection increases with a rise in the number of time points, effect size, and UMI counts. Conversely, it diminishes as the number of cells within each individual increase, along with sample-level variations (i.e., batch effects) (Additional file 2: Fig. S7). Notably, the Wilcoxon test did not fare well in all power simulations, presumably due to failure to properly control the type I error rate.

In addition, we further examined the performance of TDEseq in other two temporal expression patterns: (1) a plateau in the first few time points, then another plateau in the last few time points (we referred to this pattern as a *bi-plateau* pattern; Additional file 2: Fig. S1E), and (2) a multi-mode pattern at begin time points then stable in last time points (we referred this pattern as a *multi-modal* pattern; Additional file 2: Fig. S1F). Under the bi-plateau pattern, Mixed TDEseq still displayed more powerful performance than other methods (Additional file 2: Fig. S8A), suggesting the shape-restricted spline function is flexible to capture bi-plateau patterns. In contrast, under the multi-modal pattern, all methods achieved low detection power, but edgeR and DESeq2 showed a higher performance than other methods (Additional file 2: Fig. S8B). However, this may not be a great issue since the multi-modal pattern may be a rare scenario in real data applications [25].

### TDEseq coupled with batch removal strategy exhibits excellent performance in analyzing large heterogeneous scRNA-seq data

Intuitively, in situations with minimal or no sample-level variations (i.e., batch effects), it is reasonable to expect that trajectory-based differential expression methods (e.g., tradeSeq) would yield comparable results to temporal-based differential expression methods. To do so, we reduced the batch effect size to zero. As a result, we observed that both versions of TDEseq and tradeSeq generated well-calibrated *p*-values under the null simulations, whereas ImpulseDE2 demonstrated overly conservative *p*-values and the Wilcoxon test displays inflated *p*-values (Additional file 2: Fig. S9A). Again, both versions of TDEseq and all other approaches generated comparable results of temporal expression pattern (Additional file 2: Fig. S9B). As we know, the presence of unwanted batch effects poses substantial obstacles in detecting temporal expression changes. We therefore increased the batch effect size to 0.12. As a result, we found Mixed TDEseq outperformed other methods in terms of temporal expression pattern detection power (Fig. 1F). However, the *p*-values generated by Mixed TDEseq were not well-calibrated (Fig. 1E).

To this end, to properly control the unwanted variables in the large batch effects scenario, we additionally carried out the batch effects correction procedure prior to performing temporal gene expression analysis. To do so, we benchmarked five existing batch removal methods that can return the corrected gene expression matrix, including MNN [52], scMerge [53], ZINB-WaVE [54], ComBat [55], and Limma [56]. As a result, with evaluation criterion iLISI score [57] (Materials and Methods) for batch correction approaches, we found scMerge (0.49) achieved a higher alignment score than Limma (0.48), ComBat (0.48), MNN (0.11), and ZINB-WaVE (0.21; Additional file 2: Fig. S10). Moreover, we found Mixed TDEseq coupled with scMerge (Mixed TDEseq + scMerge) performed reasonably well in terms of the type I error control (Fig. 1E and Additional file 2: Fig. S11A) and was more powerful in detecting temporal expression genes (Fig. 1F and Additional file 2: Fig. S11B), suggesting this combination is suitable for time-course scRNA-seq data with strong sample-level variations (i.e., batch effects). Taken together, TDEseq coupled with scMerge may be an ideal approach for the identification of temporal gene expression patterns when time-course scRNA-seq data involves large heterogeneous samples.

### TDEseq performs well in the intertwined cells among time points

The simulations above all display the time point-specific expression. To mimic the cell differentiation scenario where the same type of cells were intertwined among time points (Additional file 2: Fig. S12A), we simulated additional scRNA-seq data sets (denoted as smudged data) using the *Symsim* R package [58] (Materials and Methods). Consequently, the pseudotime was inferred using Slingshot [6] according to the recommendation from the previous studies [31, 32]. In this simulation, we first took the inferred pseudotime as inputs in tradeSeq and ImpulseDE2, while the time points as inputs in both versions of TDEseq, edgeR, and DESeq2. As a result, we observed that the performance of Linear TDEseq was comparable with ImpluseDE2 in a small proportion of intertwined cells between time points (Additional file 2: Fig. S12B). With a medium proportion of intertwined cells (Additional file 2: Fig. S12C) and a large proportion of intertwined cells (Additional file 2: Fig. S12D), the pseudotime-based methods tradeSeq and ImpulseDE2 outperformed the time points-based methods, both versions of TDEseq, edgeR, and DESeq2. Furthermore, we took the inferred pseudotime as inputs in both versions of TDEseq, Linear TDEseq outperformed Mixed TDEseq, and ImpulseDE2, but not tradeSeq (Additional file 2: Fig. S12E).

In addition, we further examined the temporal expression patterns that were detected by Linear TDEseq. As a result, we found even though the pseudotime as inputs, TDEseq displayed four distinct temporal expression patterns (Additional file 2: Fig. S12F). Therefore, TDEseq was also useful for detecting temporal expression patterns with the pseudotime as inputs.

### TDEseq detects drug-associated temporal expression changes of time-course scRNA-seq data

We first applied TDEseq on a drug-treatment time-resolved scRNA-seq data set (Additional file 3: Table S1). The data were assayed by Well-TEMP-seq protocols to profile the transcriptional dynamics of colorectal cancer cells exposed to 5-AZA-CdR [23] (Materials and Methods). This scRNA-seq data set consists of D0, D1, D2, and D3 four time points (Fig. 2A), and each time point contains 4,000 cells. We expected these scRNA-seq data sets to exhibit minimal individual heterogeneity across multiple time points since Well-TEMP-seq addressed the cell lines within one chip [23] (Additional file 2: Fig. S13A). Therefore, we performed the temporal gene detection methods without batch effects correction. Since only one sample was involved in each time point, we excluded Mixed TDEseq, edgeR, and DESeq2 in this application.

**Fig. 2.**
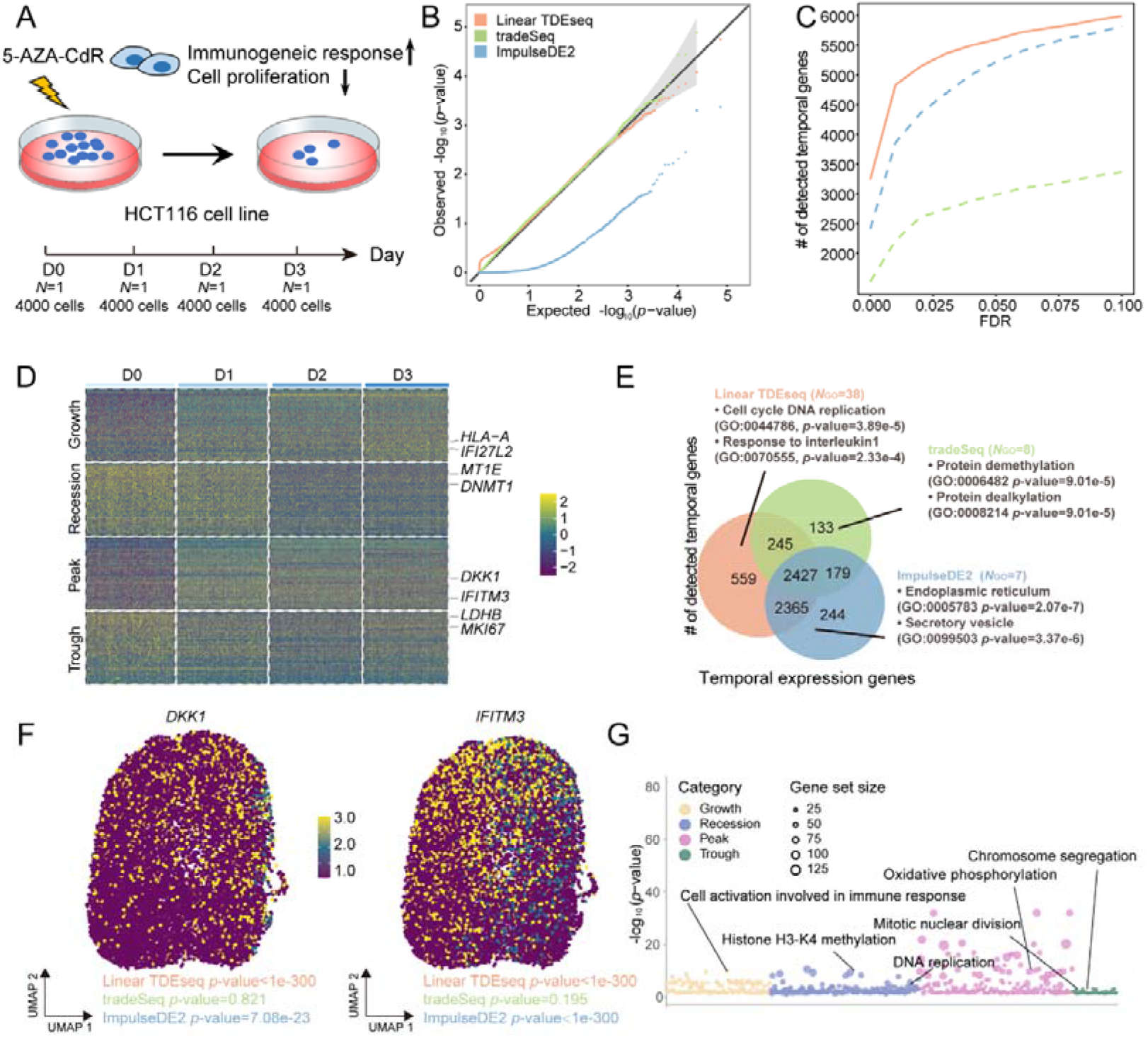
The time-resolved scRNA-seq data analysis for the HCT116 cell lines after 5-AZA-CdR treatment. (**A**) The experimental design of HCT116 cell lines treated with 5-AZA-CdR. The scRNA-seq data were assayed by Well-TEMP-seq protocols, consisting of four time points, i.e., D0, D1, D2, or D3 after treatment. (**B**) The quantile-quantile (QQ) plot shows the type I error control under the permutation strategy. The well-calibrated *p*-values will be expected laid on the diagonal line. The *p-*values produced by Linear TDEseq (orange) and tradeSeq (green) are reasonably well-calibrated, while those from ImpulseDE2 (blue) are overly conservative. (**C**) The power comparison of temporal expression gene detection across a range of FDR cutoffs. Linear TDEseq was highlighted using solid lines, while other methods were represented by dashed lines in the plots. Linear TDEseq displays the powerful performance of temporal expression gene detection. (**D**) The heatmap demonstrates the pattern-specific temporal expression genes that were identified by Linear TDEseq. Gene expression levels were log-transformed and were standardized using z-scores for visualization. The top-ranked temporal expression genes identified by Linear TDEseq show distinct four patterns. (**E**) The Venn diagram shows the overlapping of the temporally expressed genes (FDR ≤ 0.05) identified by Linear TDEseq, tradeSeq, or ImpulseDE2. Those method-specific unique genes were enriched in the number of GO terms (*N*_GO_, BH-adjusted *p*-value < 0.05). The temporal expression genes detected by Linear TDEseq were enriched with a greater number of GO terms. (**F**) The UMAP shows two temporal expression genes, i.e., *DKK1* and *IFITM3,* which were identified by Linear TDEseq but not by tradeSeq. (**G**) The Bubble plot demonstrates the significant GO terms enriched by pattern-specific temporal expression genes, which were identified by Linear TDEseq. The peak-specific temporal expression genes enriched more significant GO terms. The Wilcoxon test was excluded from this comparison due to its poor performance in simulations. DESeq2 and edgeR were excluded from this comparison due to only one sample at each time point. FDR denotes the false discovery rate.

Next, we examined the ability of Linear TDEseq in terms of type I error control. To do so, we utilized a permutation strategy (repeated 5 times) to construct a null distribution (Materials and Methods). Consistent with simulation studies, we found Linear TDEseq can produce well-calibrated *p*-values while tradeSeq produced inflated *p*-values and ImpulseDE2 produced overly conserved *p*-values (Fig. 2B). Besides, in terms of temporally expressed gene detection, Linear TDEseq outperformed other methods across a range of FDR cutoffs (Fig. 2C), even in pattern-specific temporal expression gene detection (Additional file 2: Fig. S13B). For example, Linear TDEseq identified a total of 5,596 temporally expressed genes at an FDR of 5%, including 1,341 growth genes, 1,177 recession genes, 225 trough genes, and 2,853 peak genes, which displayed four distinct temporal expression patterns (Fig. 2D). In contrast, ImpulseDE2 identified a total of 4,792 temporally expressed genes and tradeSeq detected a total of 2,672 temporally expressed genes (Additional file 3: Table S3). Overall, besides the 2,427 common shared temporally expressed genes detected by Linear TDEseq, tradeSeq, and ImpulseDE2 methods (Fig. 2E), a total of 559 temporally expressed genes were uniquely detected by TDEseq, which were also significantly enriched in cell cycle DNA replication (GO:0044786; BH adjusted *p*-value = 7.89e-e) and response to interleukin1 (GO:0070555; BH adjusted *p*-value = 0.031). In contrast, tradeSeq or ImpulseDE2 unique genes were not enriched in 5-AZA-CdR treatment response associated GO terms (Fig. 2E). Specifically, we found tumor suppressor genes which were a target of 5-AZA-CdR, i.e., *DKK1* [23, 59] was identified by TDEseq as top-ranked significant temporal expression genes (*p*-value < 1e-300, FDR = 0), while was not detected by tradeSeq (*p*-value = 0.82, FDR = 0.90), probably due to though this gene had clearly peak pattern, the log fold change was small enough, and difficult to detect with penalized splines; Besides, a 5-AZA-CdR response gene *IFITM3* [59, 60] was also identified by TDEseq as top-ranked significant genes (*p*-value < 1e-300, FDR = 0), but not detected by tradeSeq (*p*-value = 0.19, FDR = 0.36, Fig. 2F).

Finally, we performed gene set enrichment analysis (GSEA) on the pattern-specific temporal expression genes to examine top GO terms enriched by the given gene lists (Materials and Methods). Specifically, with an FDR of 5%, a total of 1,341 growth-specific temporal expression genes were detected by Linear TDEseq. These genes were enriched in a total of 179 GO terms. Due to 5-AZA-CdR treatment leads HCT116 cells to a viral mimicry state, and triggers the antiviral response [61], we expected a result of an immune response that drives the immune-associated genes up-regulated. Indeed, the GO terms contain many immune response terms, such as the cell activation involved in the immune response process (GO: 0002263; BH-adjusted *p*-value = 8.90e-7), indicating immune response was activated by the 5-AZA-CdR treatment; a total of 1,177 recession-specific temporal expression genes were enriched in a total of 244 GO terms, e.g., many regulations of histone methylation terms such as positive regulation of histone H3-K4 methylation (GO: 0051571; BH-adjusted *p*-value = 0.047), implying DNA methylation inhibitions and gene expression regulation were occurred after 5-AZA-CdR treatment, due to the global DNA demethylation effects of 5-AZA-CdR [62]; a total of 2,853 peak-specific temporal expression genes were enriched in a total of 249 GO terms. For example, The ATP metabolic process pathways, particularly oxidative phosphorylation (GO: 0006119; BH-adjusted p-value = 3.30e-7), are impacted by the increase of intracellular ROS and mitochondrial superoxide induced by 5-AZA-CdR. However, this effect diminishes over time [63]; Similarly, a total of 225 trough-specific temporal expression genes were enriched in a total of 89 GO terms, with a significant portion belonging to cell cycle pathways, including mitotic nuclear division (GO: 0140014; BH-adjusted p-value = 1.06e-6). These findings suggest that 5-AZA-CdR treatment may lead to the suppression of tumor cell proliferation and division [61] (Fig. 2G).

### TDEseq detects hepatic cell differentiation-associated temporal expression genes of time-course scRNA-seq data

We next applied TDEseq to a hepatoblast-to-hepatocyte transition study from the C57BL/6 and C3H embryo mice livers [38] (Materials and Methods; Additional file 3: Table S1). This scRNA-seq data set consists of 7 developmental stages from 13 samples, including E10.5 (54 cells from 1 sample), E11.5 (70 cells from 2 samples), E12.5 (41 cells from 2 samples), E13.5 (65 cells from 2 samples), E14.5 (70 cells from 2 samples), E15.5 (77 cells from 2 samples), and E17.5 (70 cells from 2 samples) [64] (Fig. 3A). Compared with the above time-resolved scRNA-seq data, this time-course scRNA-seq data set contains multiple samples at each time point, exhibiting small individual heterogeneity across all developmental stages (Additional file 2: Fig. S14A). Therefore, we carried out both versions of TDEseq that would be expected to be comparable in such a scenario and excluded edgeR and DESeq2 in this application due to one or two samples involved in each time point.

**Fig. 3.**
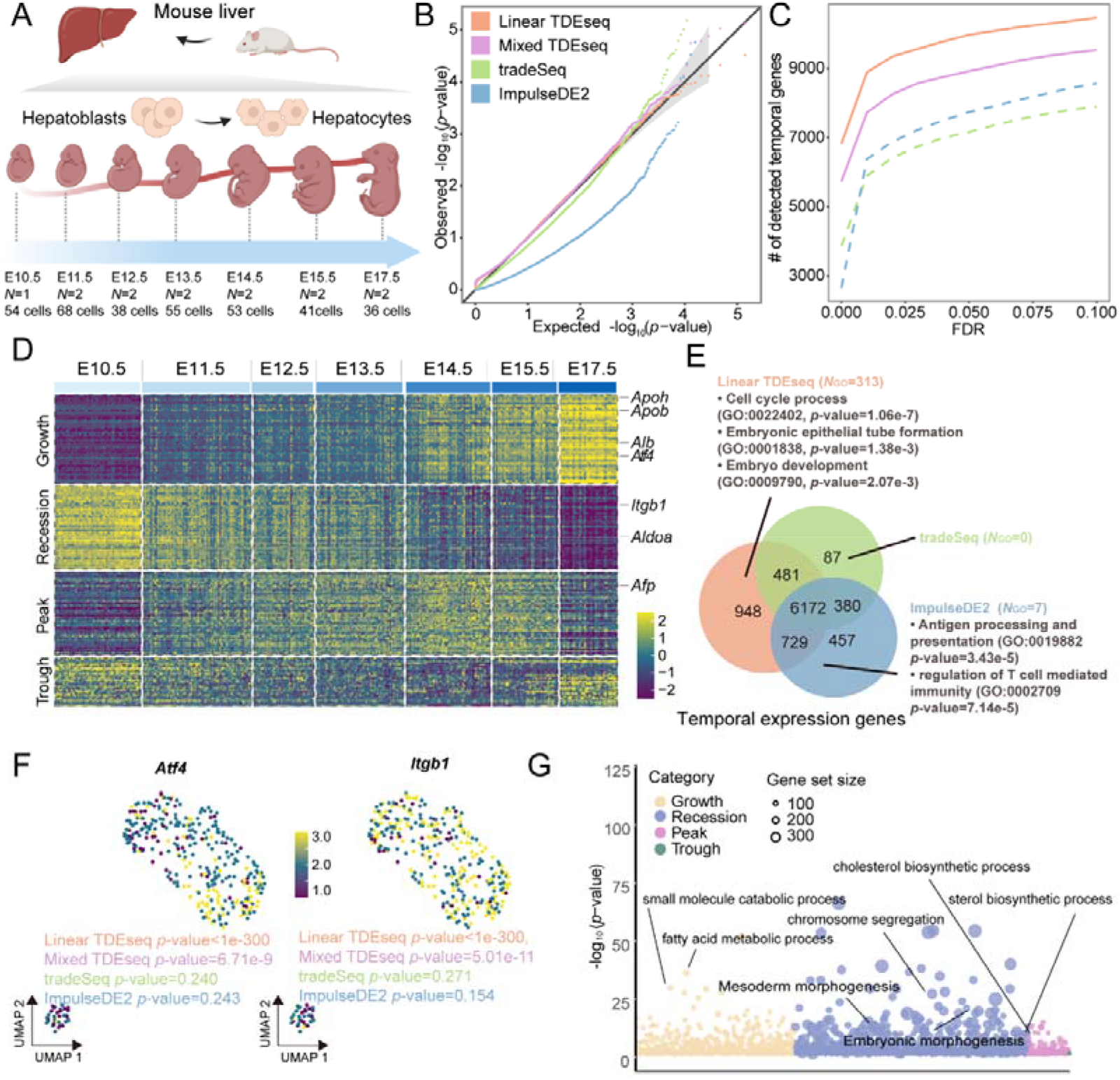
The time-course scRNA-seq data analysis for mouse fetal liver development. (**A**) The experimental design of mouse fetal liver sample collection. The scRNA-seq data were assayed on the FACS isolated cell populations, consisting of seven liver developmental stages, i.e., E10.5, E11.5, E12.5, E13.5, E14.5, E15.5, and E17.5. (**B**) The quantile-quantile (QQ) plot shows the type I error control under the permutation strategy. The well-calibrated *p*-values will be expected laid on the diagonal line. The *p-*values produced by Linear TDEseq (orange), Mixed TDEseq(plum), and tradeSeq (green) are reasonably well-calibrated, while those from ImpulseDE2 (blue) are overly conservative. (**C**) The power comparison of temporal expression gene detection across a range of FDR cutoffs. The TDEseq methods were highlighted using solid lines, while other methods were represented by dashed lines in the plots. Both versions of TDEseq display the powerful performance of temporal expression gene detection. (**D**) The heatmap demonstrates the pattern-specific temporal expression genes that were identified by Linear TDEseq. Gene expression levels were log-transformed and were standardized using z-scores for visualization. The top-ranked temporal expression genes identified by Linear TDEseq show distinct four patterns. (**E**) The Venn diagram shows the overlapping of the temporally expressed genes (FDR ≤ 0.05) identified by Linear TDEseq, tradeSeq, or ImpulseDE2. Those method-specific unique genes were enriched in the number of GO terms (*N*_GO_, BH-adjusted *p*-value < 0.05). The temporal expression genes detected by Linear TDEseq were enriched more GO terms. (**F**) The UMAP shows two temporal expression genes, i.e., *Atf4* and *Itgb1,* which were uniquely identified by Linear TDEseq. (**G**) The Bubble plot demonstrates the significant GO terms enriched by pattern-specific temporal expression genes, which were identified by Linear TDEseq. The recession-specific temporal expression genes enriched more significant GO terms, whereas trough-specific temporal expression genes were not enriched in any GO terms. The Wilcoxon test was excluded from this comparison due to its poor performance in simulations. DESeq2 and edgeR were excluded from this comparison due to only one or two samples at each time point. FDR denotes the false discovery rate.

To do so, we first examined the ability of TDEseq in terms of type I error control using permutation strategies (Materials and Methods). Consistent with the simulation results, Linear TDEseq, Mixed TDEseq, and tradeSeq could produce the well-calibrated *p*-values whereas ImpulseDE2 generated overly conservative *p*-values (Fig. 3B). Besides, in terms of temporally expressed gene detection, both versions of TDEseq outperformed other methods across a range of FDR cutoffs (Fig. 3C and Additional file 2: Fig. S14B). Specifically, Linear TDEseq identified a total of 9,975 temporally expressed genes at an FDR of 5%, including 1,266 growth genes, 7,146 recession genes, 217 trough genes, and 1,346 peak genes, which displayed four temporal distinct patterns (Fig. 3D); Mixed TDEseq identified a total of 8,924 temporally expressed genes, including 1,242 growth genes, 6,708 recession genes, 136 trough genes, and 8,38 peak genes. In contrast, ImpulseDE2 detected a total of 7,737 temporally expressed genes while tradeSeq detected a total of 7,108 temporally expressed genes (Additional file 3: Table S4). Notably, comparing with tradeSeq and ImpulseDE2, there were a total of 948 temporally expressed genes uniquely detected by Linear TDEseq at an FDR of 5% and a total of 3,517 temporally expressed genes uniquely detected by Mixed TDEseq at an FDR of 5%. Comparing the results of Linear TDEseq and Mixed TDEseq, we found the *p*-values generated from both Mixed TDEseq and Linear TDEseq demonstrated a high correlation (Spearman *R* = 0.954; Additional file 2: Fig. S14C). We further observed that some of the genes from Linear TDEseq displayed a smaller *p*-value than that from Mixed TDEseq. This observation was presumably due to Linear TDEseq being more sensitive in large sample-level variations across time points. For example, the *p*-value of a recession-specific gene *MAPK13* generated by Linear TDEseq (*p*-value = 1.0e-300) was extremely small than Mixed TDEseq (*p*-value = 1.8e-2; Additional file 2: Fig. S14D).

Moreover, the temporally expressed genes detected by both versions of TDEseq but not detected by tradeSeq or ImpulseDE2 were highly related to hepatic cell differentiation (Fig. 3E), where many of them have been validated by the previous studies [38, 65, 66]. For example, we found a key mouse fetal liver development regulator *Atf4* [66], which exhibits a growth pattern (Fig. 3F), was only identified by TDEseq as the top-ranked significant temporal expression gene (*p*-value < 1e-300, FDR = 0), while was not detected by tradeSeq (*p*-value = 0.240, FDR = 0.275) and ImpulseDE2 (*p*-value = 0.243, FDR = 0.079). Besides, *Itgb1* displays the growth pattern (bi-plateau pattern; Fig. 3F) for liver microstructure establishment during the embryonic process, which was only identified by TDEseq as the top-ranked significant temporal expression gene (*p*-value < 1e-300, FDR = 0), while was not detected by tradeSeq (*p*-value = 0.272, FDR = 0.309). These genes uniquely detected by Linear TDEseq were enriched in the liver embryo process, particularly the cell cycle process (GO:0022402, BH-adjusted *p*-value=1.33e-5) and embryo development (GO:0009790; BH-adjusted *p*-value = 0.0361); whereas the temporal expression genes uniquely detected by tradeSeq or ImpulseDE2 were not enriched in liver development-related gene sets (Fig. 3E), and ImpulseDE2 wrongly detected the peak or trough pattern genes as growth pattern genes (Additional file 2: Fig. S14E). In addition, the enrichment analysis of unique temporal expression genes from Mixed TDEseq and Linear TDEseq showed similar results (Additional file 2: Fig. S14F).

Next, we performed GSEA on the pattern-specific temporal expression genes identified by Linear TDEseq, to examine the four pattern-specific functions during hepatic cell differentiation (Materials and Methods). Specifically, with an FDR of 5%, a total of 1266 growth-specific temporal expression genes were enriched in a total of 685 GO terms. Notably, these growth-related genes were significantly enriched in liver function-associated pathways, reflecting the mature process from hepatoblasts to hepatocytes. For example, almost all of these enriched terms were associated with metabolic processes, biosynthetic processes, or organic substance transport, with key functions attributed to mature hepatocytes (Fig. 3G). Particularly, the fatty acid metabolic process (GO:0006631; BH-adjusted *p*-value = 4.99e-37), lipid catabolic process (GO:0016042; BH-adjusted *p*-value = 3.34e-26), and secondary alcohol metabolic process (GO:1902652; BH-adjusted *p*-value = 8.06e-16), which are the main function of mature hepatocytes. On the other hand, a total of 7,146 recession-specific temporal expression genes were enriched in 1,000 GO terms. Interestingly, these GO terms were related to embryo or tissue development (Fig. 3G), such as embryonic morphogenesis (GO:0048598; BH-adjusted *p*-value = 1.04e-3) and mesoderm morphogenesis (GO:0048332; BH-adjusted *p*-value = 4.05e-3). This finding suggests the involvement of an embryo development process, possibly linked to organogenesis occurring at the E14.5 stage [67]. Furthermore, these genes may signify the loss of embryonic cell identity in mature hepatocytes.

Finally, since the scRNA-seq data showed intertwined cells among time points (Additional file 2: Fig. S14G), we further applied TDEseq with the pseudotime as inputs. As a result, we observed both versions of TDEseq generated comparable results with tradeSeq and ImpulseDE2 for temporal expression gene detection (Additional file 2: Fig. S14H). Besides, these genes demonstrated distinct temporal expression patterns for TDEseq (Additional file 2: Fig. S14I).

Taken together, we found both versions of TDEseq yields similar results in terms of type I control rate and temporal expression gene detection when scRNA-seq data exhibits small individual heterogeneity over time points. Therefore, considering the computation burden for large-scale scRNA-seq data applications, we recommended Linear TDEseq in a small individual heterogeneity scenario.

### TDEseq detects the epithelial cell evaluation-associated temporal expression genes of time-course scRNA-seq data

We again applied TDEseq to detect temporal expression genes altered in human metastatic lung adenocarcinoma (LUAD) cancer [14] (Materials and Methods). Here, we were primarily interested in epithelial cells of this time-course scRNA-seq data, which involves a total of five distinct evolution stages, i.e., stage normal, stage I, stage II, and stage III and stage IV (Additional file 3: Table S1). Since stage II contains a relatively small number of cells (i.e., 119 cells) compared with other stages, we excluded this stage, resulting in a total of 3,703 cells from 11 samples in the normal stage, 5,651 cells from 8 samples in stage I, 1,500 cells from 2 samples in stage III, and 3,053 cells from 7 samples in stage IV (Fig. 4A). We noticed that these scRNA-seq data sets contain sample-level variations across stages (iLISI = 0.10; Additional file 2: Fig. S15A). Therefore, we performed both versions of TDEseq in such a scenario.

**Fig. 4.**
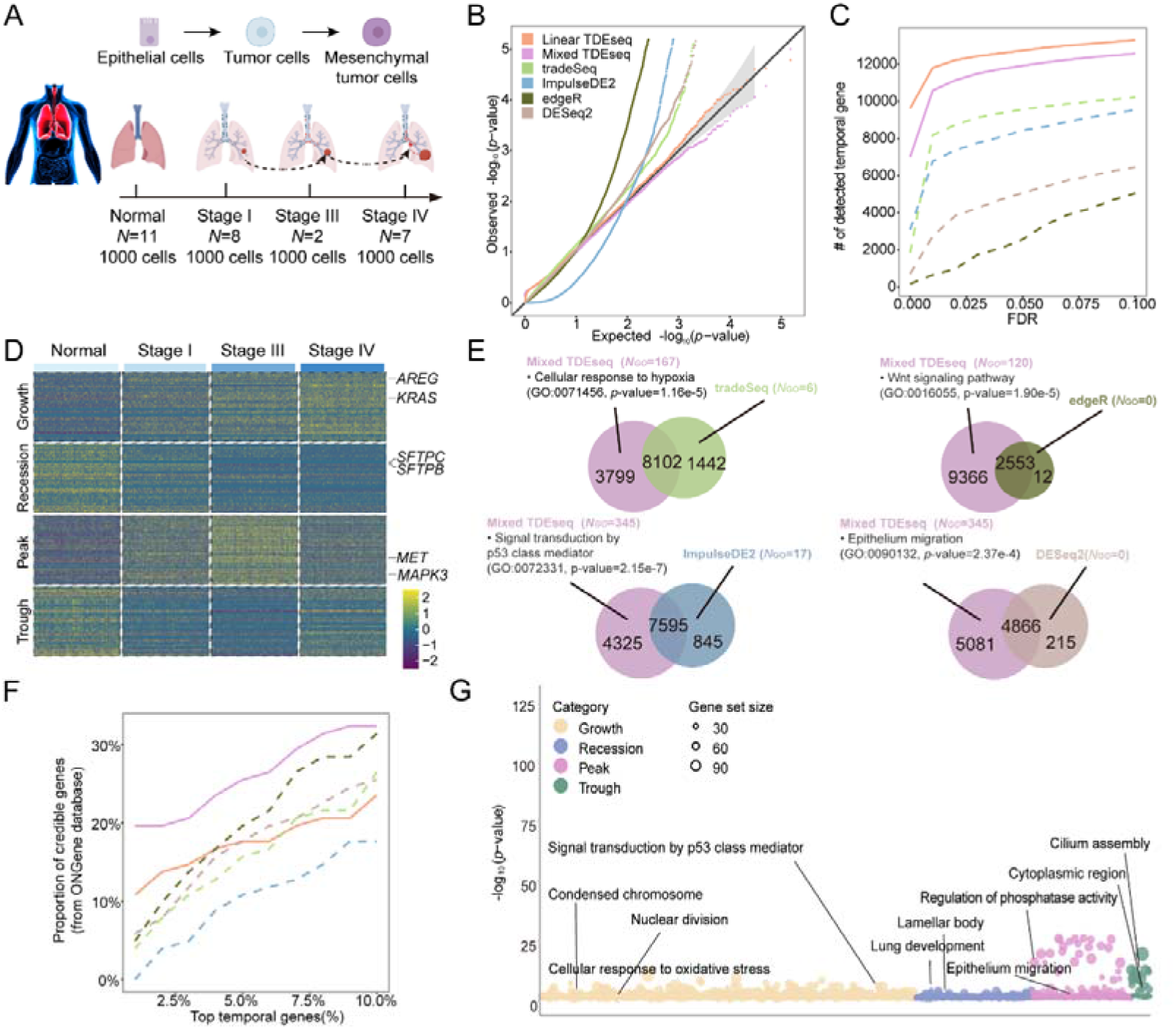
The time-course scRNA-seq data analysis for human metastatic LUAD. (**A**) The experimental design of human lung sample collection. The scRNA-seq data were assayed by 10X Genomics Chromium protocols, consisting of 4 LUAD evaluation stages, i.e., Normal, Stage I (early LUAD), Stage III (advanced LUAD), and Stage IV (lymph node metastasis). (**B**) The quantile-quantile (QQ) plot shows the type I error control under the permutation strategy. The well-calibrated *p*-values will be expected laid on the diagonal line. The *p-*values produced by Linear TDEseq (orange), and Mixed TDEseq(plum) are reasonably well-calibrated, while those from tradeSeq (green), ImpulseDE2 (blue) edgeR (dark green), and DESeq2 (brown) are inflated. (**C**) The power comparison of temporal expression gene detection across a range of FDR cutoffs. The TDEseq methods were highlighted using solid lines, while other methods were represented by dashed lines in the plots. Both versions of TDEseq display the powerful performance of temporal expression gene detection. (**D**) The heatmap demonstrates the pattern-specific temporal expression genes that were identified by Mixed TDEseq. Gene expression levels were log-transformed and were standardized using z-scores for visualization. The top-ranked temporal expression genes identified by Mixed TDEseq show distinct four patterns. (**E**) The Venn diagram shows the overlapping of the temporally expressed genes (FDR ≤ 0.05) in pairwise comparisons between Mixed TDEseq and tradeSeq, ImpulseDE2, DESeq2, and edgeR. Those method-specific unique genes were enriched in the number of GO terms (*N*_GO_, BH-adjusted *p*-value < 0.05). Many more GO terms were enriched in the Mixed TDEseq-unique temporal expression genes than in other methods. (**F**) The proportion of enrichment for the detected temporal expression genes. The given gene set (136 genes) was collected from ONGene [71] database. Mixed TDEseq enriched more temporal genes than other methods across a range of top-number cutoffs. (**G**) The Bubble plot demonstrates the significant GO terms enriched by pattern-specific temporal expression genes, which were identified by Mixed TDEseq. The Wilcoxon test was excluded from this comparison due to its poor performance in simulations. FDR denotes the false discovery rate.

To do so, we first examined the ability of temporal gene detection methods in terms of type I errors. As we expected, when large sample-level variations were involved, Mixed TDEseq and Linear TDEseq produced well-calibrated *p-*values. The other methods tradeSeq, ImpulseDE2, DESeq2, and edgeR generated the inflated *p-*values (Fig. 4B). Either in terms of temporal expression gene detection (Fig. 4C) or in terms of temporal expression pattern detection (Additional file 2: Fig. S15B), both versions of TDEseq outperformed other methods across a range of FDR cutoffs. Specifically, Mixed TDEseq identified a total of 11,919 temporally expressed genes at an FDR of 5%, which displayed four temporal distinct patterns (Fig. 4D), while Linear TDEseq detected 12,263 temporal genes, tradeSeq detected 9,562 temporal genes, ImpulseDE2 identified 8,440 temporal genes, DESeq2 detected 5,081 temporal genes and edgeR detected 2,565 temporal genes (Additional file 3: Table S5).

To validate whether the temporal expression genes were related to epithelial cell evaluation, we performed the following two lines of enrichment analyses. We examined the temporal expression genes detected by Mixed TDEseq but not detected by tradeSeq, ImpulseDE2, DESeq2, or edgeR, finding many genes were associated with LUAD evolution. For example, a LUAD driver *MAP2K1* [68] was detected by Mixed TDEseq as significant temporal expression genes that gradually up-regulated during LUAD progression (*p*-value = 1.17e-7, FDR = 0), while not detected by ImpulseDE2 (*p*-value = 0.553, FDR = 0.105); Besides, another LUAD drivers *KRAS* [68] was also detected by Mixed TDEseq as significant temporal expression genes that gradually up-regulated during LUAD progression (*p*-value = 4.05e-12, FDR = 0), while not detected by tradeSeq (*p*-value = 0.089, FDR = 0.356). Furthermore, we performed pairwise comparisons of Mixed TDEseq vs other methods. The result shows the unique temporal expression genes from Mixed TDEseq were enriched in GO terms that related to the LUAD progression (Fig. 4E), such as signal transduction by p53 class mediator [69] (GO:0072331; BH-adjusted *p*-value = 1.09e-4) and Cellular response to hypoxia [70] (GO:00071456; BH-adjusted *p*-value = 4.91e-3). On the other hand, we curated a total of 136 LUAD-related genes from the ONGene database [71] (Additional file 3: Table S6) to highlight the importance of temporal expression genes detected by different methods. As a result, we found that the temporal expression genes from Mixed TDEseq were enriched more genes than other methods across a range of top genes (Fig. 4F). Notably, though Linear TDEseq detected more temporal genes, the temporal genes uniquely identified by Mixed TDEseq were enriched in more biologically meaningful GO terms (Additional file 2: Fig. S15C), e.g., epithelial cell migration (GO:0010631; BH-adjusted *p*-value = 0.048).

Finally, we performed GSEA on the pattern-specific temporal expression genes to examine the four pattern-specific functions during LUAD progression. Specifically, with an FDR of 5%, Mixed TDEseq detected a total of 3,249 growth-specific temporal expression genes, which were enriched in 812 GO terms (Fig. 4G). The top GO terms contained many tumor proliferation or metastasis-associated pathways, such as signal transduction by p53 class mediator [69] (GO:0072331; BH-adjusted *p*-value = 1.57e-6), regulation of canonical Wnt signaling pathway [72] (GO:0060828; BH-adjusted *p*-value = 2.81e-3), suggesting epithelial cells proliferation towards tumor cells; Mixed TDEseq identified a total of 3,671 recession-specific temporal expression genes, which were enriched in 276 GO terms (Fig. 4G). Those genes would be expected enriched in normal lung function terms as a result of the process of low-grade tumors developing into high-grade tumors. Indeed, the top GO terms contained lung development (GO:0030324; BH-adjusted *p*-value = 8.26e-3) and lamellar body (GO:0042599; BH-adjusted *p*-value = 4.08e-4), suggesting the proliferation process of epithelial cells towards tumor cells. TDEseq detected a total of 2,526 peak-specific temporal genes, which were enriched in a total of 244 GO terms. Notably, those peak genes were further enriched in hypoxia pathways, such as response to oxidative stress (GO:0006979; BH-adjusted *p*-value = 1.19e-2), as well as epithelium migration (GO:0090132; BH-adjusted *p*-value = 4.56e-2). This evidence further validated the fact that hypoxia occurs in the intermediate stages of LUAD promoting lymphatic metastasis [70].

Taken together, Mixed TDEseq can address time-course scRNA-seq data with relatively large sample-level variations over time points. However, one of the concerns regarding whether batch effects removal can improve the identification of temporal expression genes. To do so, we performed the temporal gene detection analysis using Mixed TDEseq either with or without batch correction. We found that Mixed TDEseq can generate well-calibrated *p*-values in both cases (Additional file 2: Fig. S15D). In terms of temporal expression gene detection, Mixed TDEseq alone would produce a more powerful performance than that with batch correction across a range of FDR cutoffs (Additional file 2: Fig. S15E). The slightly poor performance of Mixed TDEseq coupled with scMerge may be due to the over-correction of sample-level variations. Furthermore, we observed the temporal expression genes uniquely identified by Mixed TDEseq were significantly enriched in LUAD-related pathways (Additional file 2: Fig. S15F), suggesting Mixed TDEseq well-addressed time-course scRNA-seq data with relatively large sample-level variations.

### TDEseq detects NK cell response temporal genes of time-course scRNA-seq data

We finally applied TDEseq to detect the temporal expression changes of natural killer (NK) cells from 21 severe/critical COVID-19 patients [9] (Materials and Methods). This time-course scRNA-seq data set contains 19 time points (Additional file 3: Table S1), which could be grouped into five developmental stages (Fig. 5A), i.e., stage I (consisting of 930 cells from 3 patients), stage II (939 cells from 4 patients), stage III (893 cells from 3 patients), stage IV (768 cells from 3 patients), stage V (1,000 cells from 8 patients). Since these scRNA-seq data sets contain large sample-level variations across different stages (iLISI = 0.11; Fig. 5B), presumably due to a large number of heterogeneous patients involved in this study. To do so, following the results from simulation studies, we first carried out Mixed TDEseq with or without the batch effect removal procedure using scMerge [53]. Since tradeSeq and ImpulseDE2 have originally built-in variables to control sample-level variations. For a fair comparison, we additionally incorporated the sample indicator variables as covariates in both tradeSeq and ImpulseDE2 models.

**Fig. 5.**
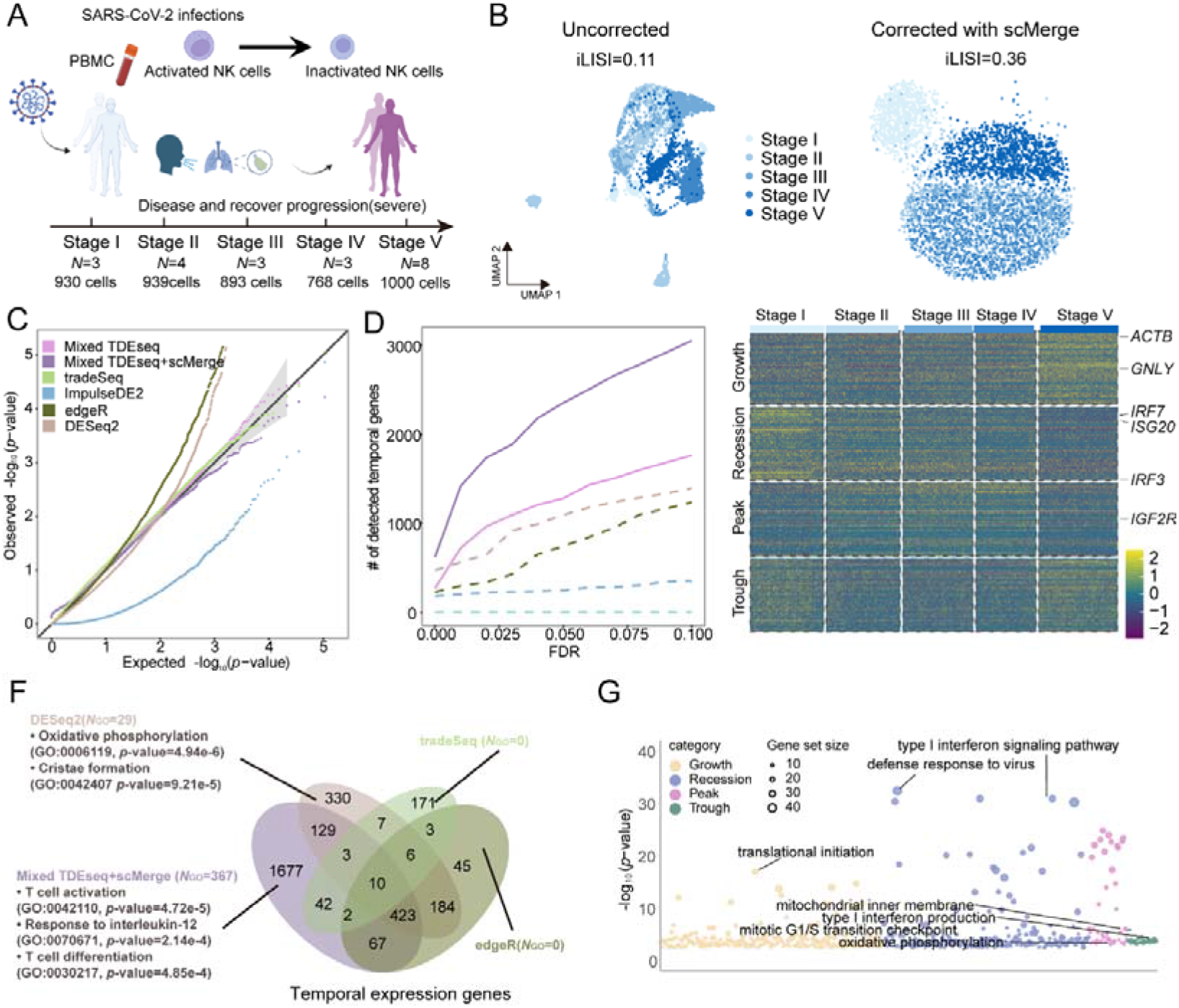
The time-course scRNA-seq data analysis for the NK cell response to SARS-COV-2 infection. (**A**) The experimental design of SARS-COV-2 infection samples from PBMC. The scRNA-seq data were assayed by 10X Genomics Chromium protocols, consisting of 5 stages, i.e., stage I (4-8 days), stage II (10-13 days), stage III (19-24 days), stage IV (28-34 days) and stage V (110-123 days). (**B**) The UMAP demonstrates cell alignment from different stages. These scRNA-seq data sets display strong batch effects over heterogeneous samples (iLISI = 0.10, left panel). The cells are well-aligned after performing integrative analysis using scMerge (iLISI = 0.36, right panel). (**C**) The quantile-quantile (QQ) plot shows the type I error control under the permutation strategy. The well-calibrated *p*-values will be expected laid on the diagonal line. The *p-*values produced by Mixed TDEseq (plum), Mixed TDEseq coupled with scMerge (purple), and tradeSeq (green) are reasonably well-calibrated, while those from ImpulseDE2 (blue) are overly conservative, and those from edgeR (dark green) and DESeq2 (brown) are inflated. (**D**) The power comparison of temporal expression gene detection across a range of FDR cutoffs. The TDEseq methods were highlighted using solid lines, while other methods were represented by dashed lines in the plots. TDEseq coupled with scMerge is more powerful in identifying more temporal expression genes than other comparative methods. (**E**) The heatmap demonstrates the pattern-specific temporal expression genes that were identified by Mixed TDEseq coupled with scMerge. Gene expression levels were log-transformed and then standardized using z-scores for visualization. The top-ranked temporal expression genes identified by Mixed TDEseq coupled with scMerge show distinct four patterns. (**F**) The Venn diagram shows the overlapping of the temporally expressed genes (FDR ≤ 0.05) identified by Mixed TDEseq coupled with scMerge, tradeSeq, DESeq2, and edgeR. ImpulseDE2 was excluded because it only identified 3 temporal DE genes. Those method-specific unique genes were enriched in the number of GO terms (*N*_GO_, BH-adjusted *p*-value < 0.05). The temporal expression genes detected by Mixed TDEseq coupled with scMerge enriched more GO terms. (**G**) The Bubble plot demonstrates the significant GO terms enriched by pattern-specific temporal expression genes, which were identified by Mixed TDEseq coupled with scMerge. The Wilcoxon test was excluded from this comparison due to its poor performance in simulations. FDR denotes the false discovery rate.

As we expected, in terms of type I error control, Mixed TDEseq, and Mixed TDEseq coupled with scMerge (Mixed TDEseq+scMerge) can produce well-calibrated *p*-values (Fig. 5C). Besides, in terms of temporal expression gene detection, Mixed TDEseq coupled with scMerge detected more temporal expression genes than Mixed TDEseq across a range of FDR cutoffs (Fig. 5D, Additional file 3: Table S7). In addition, with a range of FDR cutoffs Mixed TDEseq+scMerge identified more pattern-specific temporal genes (Additional file 2: Fig. S16A), which displayed four temporal distinct patterns (Fig. 5E). Moreover, the temporal expression genes that were uniquely detected by Mixed TDEseq+scMerge were enriched in the defense response to the virus GO term (GO:0051607) [73] (Additional file 3: Table S8; Additional file 2: Fig. S16B). Therefore, we performed the Mixed TDEseq+scMerge in the following analysis.

Notably, many top temporal expression genes uniquely detected by Mixed TDEseq coupled with scMerge were highly related to the NK cell response to COVID-19 infection. For example, *GNLY* which highly expressed in healthy people than patients with viral infections [74] was identified by Mixed TDEseq+scMerge as a significant temporal expression gene (*p*-value = 2.93e-5, FDR = 0.0; Additional file 2: Fig. S16C), while was not detected by tradeSeq (*p*-value = 0.377, FDR = 1.0), ImpulseDE2 (*p*-value = 0.994, FDR = 1), DESeq2 (*p*-value = 0.028, FDR = 0.051) or edgeR (*p*-value = 0.015, FDR = 0.093). Another example is the *ILF3* gene which plays an important role in the establishment of type I IFN antiviral program[75], was uniquely identified by Mixed TDEseq+scMerge (*p*-value = 2.67e-4, FDR = 1.16e-3; Additional file 2: Fig. S16C) but did not detect by tradeSeq (*p*-value = 0.746, FDR = 1.0), ImpulseDE2 (*p*-value = 0.952, FDR = 1.0), DESeq2 (*p*-value = 0.684, FDR = 0.712) or edgeR (*p*-value = 0.904, FDR = 0.892). This evidence supported that the temporal DE genes detected by Mixed TDEseq+scMerge were more specific to the SARS-COV-2 response biological process. Consequently, we further performed the GSEA on the temporal expression genes uniquely detected by Mixed TDEseq+scMerge, enriched in the immune response to virus infection pathways such as T cell activation (GO:0042110; BH-adjusted *p*-value = 0.027), and response to interleukin-12 (GO:0070671; BH-adjusted *p*-value = 0.029) [76] (Fig. 5F).

Finally, we performed GSEA on the pattern-specific temporal expression genes. Specifically, with an FDR of 5%, Mixed TDEseq+scMerge detected a total of 654 growth-specific temporal expression genes, which were significantly enriched in cell cycle-associated pathways, such as mitotic G1/S transition checkpoint (GO:0044819; BH-adjusted *p*-value = 7.81e-3). It was shown that NK cells showcased up-regulated patterns of cell cycle and division after SARS-COV-2 infection [77]; also enriched in the cellular response to interleukin-12 (GO:0071349; BH-adjusted *p*-value = 1.33e-2), because IL-12 promotes NK cell proliferation at the end stage of SARS-COV-2 infection [78]; Mixed TDEseq+scMerge detected a total of 809 recession-specific temporal expression genes, which were enriched in the immune response to the virus as a result of SARS-COV-2 infection. Indeed, most of the top GO terms were immune response-associated pathways, such as defense response to the virus (GO:0051607; BH-adjusted *p*-value = 1.76e-29) and type I interferon signaling pathway (GO:0060337; BH-adjusted *p*-value = 1.81e-28) which promotes NK cell expansion during viral infection [79]; Mixed TDEseq+scMerge detected a total of 567 peak-specific temporal expression genes, which were enriched in oxidative phosphorylation (GO:0006119; BH-adjusted *p*-value = 0.025) and mitochondrial gene expression (GO:0140053; BH-adjusted *p*-value *=* 4.11e-2), consistent with that long period of activation enhances effector functions in the NK cells and upregulated OXPHOS [80, 81] (Fig. 5G).

Taken together, we found Mixed TDEseq coupled with scMerge performed effectively in mitigating substantial sample-level variations (i.e., batch effects), which were presented in time-course scRNA-seq data. However, the batch correction methods do introduce extra variations into the data. Therefore, a more comprehensive assessment of the performance of temporal gene testing is required to determine whether these variations are advantageous, particularly for sparse scRNA-seq data [29].

## DISCUSSION

In this paper, we have presented TDEseq, a non-parametric statistical method designed for identifying temporal expression patterns in time-course single-cell RNA sequencing (scRNA-seq) data. By incorporating shape-constrained spline models, TDEseq typically enables the detection of four specific temporal patterns, i.e., growth, recession, peak, or trough. Two versions of TDEseq (i.e., *Mixed TDEseq* and *Linear TDEseq*) were developed to accommodate different real data application scenarios. Specifically, Mixed TDEseq was designed for analyzing the time-course scRNA-seq data with heterogeneous samples and large sample-level variations (i.e., batch effects) across time points, such as cancer development, while Linear TDEseq was tailored for handling the data with small heterogeneous samples, such as cell differentiation. With extensive simulations and four real data applications, TDEseq generated well-calibrated *p*-values and demonstrated powerful detection of temporal expression genes, highlighting its robustness and reliability in time-course scRNA-seq data analysis.

Particularly, the statistical modeling of TDEseq is different from tradeSeq and ImpluseDE2. TDEseq incorporates either *I*-splines [35] or *C*-splines [36] to model temporal expression patterns, and builds upon linear additive mixed models (Materials and Methods) to characterize the dependent cells within an individual. TDEseq was originally designed for time-course scRNA-seq studies, in which cells for each time point were not largely intertwined between adjacent time points. In contrast, tradeSeq [31] employs a generalized additive model framework to model gene expression profiles of pseudotime for different lineages; while ImpluseDE2 [25] relies on a descriptive impulse function [82] to distinguish permanently from transiently up- or down-regulated genes over multiple time points. As a result, we observed that the *p*-values from tradeSeq and ImpluseDE2 under permuted null were not well-calibrated even to control the sample-level variations as covariates, presumably due to its inability to model dependent cells within an individual rather than independent cells. We also found that the power of tradeSeq or ImpluseDE2 was lower than that of TDEseq (Additional file 2: Fig. S7), likely due to its suitability for detecting a distinct type of differential expression patterns along a lineage or between lineages. In addition, we also developed a linear version of TDEseq to ensure more scalable computation for large-scale scRNA-seq data but with small batch effects. As a result, we found the performance of Linear TDEseq was comparable with Mixed TDEseq in small sample heterogeneity (Fig. 1C and 1F). Besides, we found two pseudo-bulk aggregation methods, DESeq2, and edgeR, performed well in both high UMI counts and small number of cells per sample scenarios (Additional file 2: Fig. S3D and S4C). However, the pseudo-bulk aggregation methods potentially generated biased inference and underpowered results due to the cells from the same individual are not statistically independent [83]. Finally, we found TDEseq demonstrates powerful performance in capturing the temporal expression patterns with the intertwined nature of cells across time points, particularly in the scenarios characterized by a modest proportion of intertwined cells between time points. Nevertheless, with the case where a substantial proportion of cells exhibit intertwined across time points, we suggest the pseudotime-based methods are used for detecting temporal expression genes. Exploration of the extent of cell intertwining could be pursued through trajectory inference or leveraging existing biological insights.

Based on the aforementioned observations, the sample-level variation (i.e., batch effect) plays a crucial role in the identification of temporal expression genes. However, the removal of this unwanted variation in time-course scRNA-seq data poses significant challenges [84, 85], and there are currently no efficient criteria to measure these effects. Fortunately, we provide a solution called TDEseq, which can effectively handle such unwanted variation across multiple time points in time-course scRNA-seq studies. In contrast, tradeSeq and ImpluseDE2 directly model gene expression raw counts and incorporate the covariates to control the individual heterogeneity. However, this approach leads to a significant loss in power for detecting temporal expression genes and inevitably increases the computational burden. This is a reason why TDEseq was designed to model the transformed gene expression level (e.g., log-normalized or variance stabilizing transformation) rather than the count nature of raw gene expression data, allowing for scRNA-seq data preprocessing prior to performing TDEseq. Furthermore, even in the presence of large individual heterogeneity in scRNA-seq data, TDEseq, when coupled with batch removal methods, offers a promising approach to identifying temporal expression genes. In this study, we evaluated five batch effects removal methods. The results indicated that scMerge displays a more powerful performance than the other four methods. However, it is worth noting that the simulated scRNA-seq data may not fully mimic real-time-course scRNA-seq data. Therefore, alternative batch removal methods could also be applied to TDEseq analysis, especially a well-designed batch correction method specifically tailored for time-course scRNA-seq data, which would greatly enhance the performance of TDEseq.

Finally, several potential extensions for TDEseq can enhance its capabilities. Presently, TDEseq is designed to identify four distinct patterns—growth, recession, peak, and trough. In our efforts to broaden the scope of temporal expression patterns, we explored the bi-plateau pattern. While TDEseq demonstrated powerful performance in this pattern, it faced challenges in handling multi-modal temporal expression patterns. Besides, we observed that Mixed TDEseq detected weak peak or weak trough patterns as growth or recession patterns; while Linear TDEseq detected weak peak or weak trough patterns as peak or trough patterns making the difficult determination of temporal expression patterns. This would be a meaningful exploration in future research. Furthermore, the parameter inference of LAMM, the underlying model of TDEseq, becomes notably challenging when the number of cells is large (number of cells > 6,000). To address this issue, it would be beneficial to incorporate more efficient algorithms that reduce the computation burden. Alternatively, a down-sampling strategy or a pseudo-cell [2, 86] strategy could be employed for each time point when dealing with an extremely large number of cells.

Overall, TDEseq is well-suited to analyzing time-course scRNA-seq data sets. Thus, it can be flexibly deployed to investigate the important temporal expression genes and their potential roles during growth, development, or disease progression.

## CONCLUSION

In this paper, we present an algorithm TDEseq for the identification of temporal expression genes in time-course scRNA-seq data. To detect the temporal expression genes, we propose a linear additive mixed model that relies on shape-constrained spline. TDEseq accounts for the correlated nature of cells within individuals and demonstrates robustness in the presence of cellular heterogeneity and sample variations. With and extensive and comprehensive evaluation on various data sets, including both synthetic and real scRNA-seq data, TDEseq has shown superior performance against other methods such as tradeSeq, ImpulseDE2, Wilcoxon test, DESeq2 and edgeR. Overall, TDEseq stand as a powerful tool enabling the precise identification of temporal expression genes in time-course scRNA-seq data and facilitating a deeper understanding of the dynamic biological processes.

## MATERIALS AND METHODS

### Models and Algorithm

As the aforementioned overview, TDEseq typically models the log-normalized gene expression levels along the multiple time points as inputs. Subsequently, TDEseq was performed to identify temporal expression genes that display one of four possible patterns (i.e., growth, recession, peak, or trough). Specifically, we assume the transformed gene expression level *y_gji_*(*t*) for gene *g*, individual *j* and cell *i*, at time point *t* is,

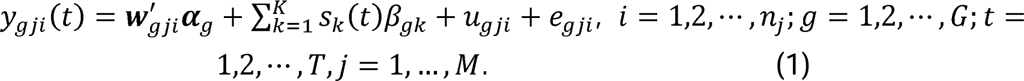

where ***w**_gji_* is the cell-level or time-level covariate (e.g., cell size, or sequencing read depth), ***α**_g_* is its corresponding coefficient; ***u**_g_* is a random vector to account for the variations from heterogeneous samples, i.e.,

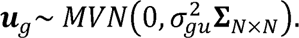

where **Σ**_*N×N*_ is a block diagonal matrix with a total of *M* block matrices, in which all elements of **Σ**_*n_j_×n_j_*_ are ones; *n_j_* is the number of cells for the individual or replicate *j* and 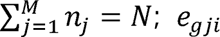 is a random effect, which is an independent and identically distributed variable that follows a normal distribution with mean zero and variance 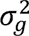 to account for independent noise, i.e.,

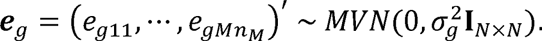

where *s_k_* is a smoothing spline basis function to characterize the temporal gene expression patterns. The regression function is estimated by a linear combination of the basis function with constrained. In particular, the *I*-splines defined as 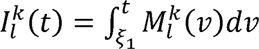 [35] were used to characterize both growth and recession patterns; while *C*-splines as 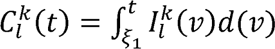 [36], taking integral operation of *l*-splines, were used to characterize both peak and trough patterns, where the order 1 *M*-splines are computed as,

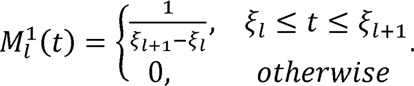

The order *k M*-splines are computed as

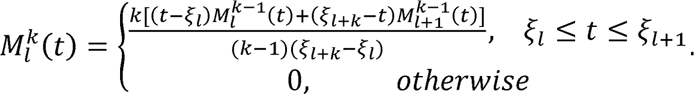

where the number of *M*-splines basis functions is *k* + *l*; *l* is the number of grid points; *k* is the order of splines; we define knots 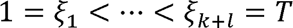. With the spline functions *s*_1_(*t*), ..., *s_K_(t)*, we infer the parameters of Equation 1 for gene *g*, which could be translated into estimating the parameters of the equation,

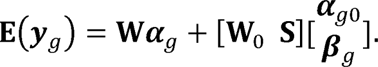

where ***W***_0_ is the linear part of the spline basis function and ***S*** is the nonlinear pacrt:of the scp:line basis function which is an *N* × (*k* + *l*) matrix with columns *s*_1_(*t*), ... *s*_*k+l*_(*t*) is the number of knots of the spline functions; For the growth or recession patterns, **S**, was assigned by *l*-spline basis functions and **W**_0_ = [**1**_*N*_] For both peak and trough patterns, **S** was assigned by the *C*-spline basis vectors and **W**_0_ = [**1**_*N*_, *t*], where ***t*** = (*t*_1_,...,*t_N_*’ represents the time points. **1**_*N*_ is an all-ones vector; **W** is a covariates matrix; ***α***_*g*_, ***α***_*g*0_ and ***β**_g_* are the corresponding regression coefficients. Hence, the complete likelihood 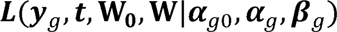 of the transformed gene expression data ***y**_g_* for gene *g* at a time, *t* is:

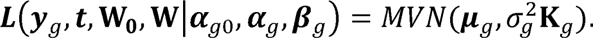

Where 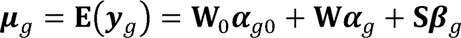 is the mean and 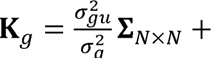 **I***_N×N_* is the covariance matrix; *MVN*(·) represents multivariate normal distribution. For any pattern constraints, we assume a linearly independent set of **S,W**_0_ and **W** together as a closed convex cone:

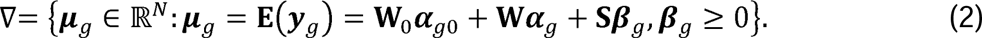

The parameter estimation of TDEseq models is notoriously difficult, as it involves the calculation of a matrix determinant and a matrix inversion of **K**_*g*_. To enable The parameter estimation of TDEseq models is notoriously difficult, as it involves scalable estimation and inference for TDEseq models, we have developed an efficient inference algorithm that: 1) performs a block diagonal matrix (one individual as an all-one block matrix) eigen-decomposition (Additional file 1: Supplementary Text), i.e., 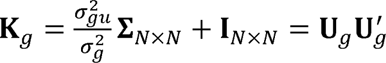; 2) transforms the variables of Equation 2 as: 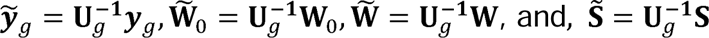 resulting in

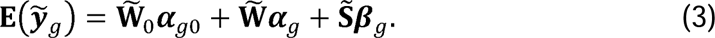

and the following new closed convex cone:

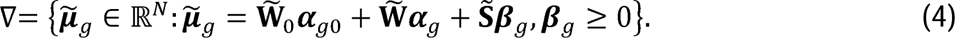

To efficiently estimate the parameter of an ordinary linear regression model (Equation 3) with cone constraint (Equation 4), we developed a cone projection algorithm following previous work [41]. With the estimated parameters, we further examined the gene expression pattern-specific parameter ***β**_g_*, which follows a mixture of *Beta* distributions [87] (Additional file 1: Supplementary Text). Finally, TDEseq returns *p*-values for four temporal expression patterns, i.e., growth, recession, peak, and trough.

### The choice of parameter knots in TDEseq models

The number of knots can significantly influence the smoothness and flexibility of the resulting curve [88]. Increasing the number of knots in general results in a more adaptable spline, enhancing its ability to capture intricate and irregular data patterns. Nonetheless, excessive flexibility in a spline can lead to overfitting, causing it to closely mimic noise in the data instead of representing the fundamental trend. Conversely, using too few knots can lead to underfitting, resulting in an overly smooth spline that misses crucial data features. Particularly, in this paper, we found that temporal gene detections are typically more robust with varying the number of knots across a range of FDR cutoffs (Additional file 2: Fig. S17). TDEseq performed well when the number of knots *k* equals the number of time points. Therefore, we set the number of knots of TDEseq as the number of time points by default.

### Linear TDEseq

Linear TDEseq is a reduced special model of Mixed TDEseq (Equation 1), which drops the random effect term *u_gji_*, typically to model only one individual involved in each stage or small individual heterogeneity across all stages in time-course scRNA-seq data. Specifically, we assume the log-normalized gene expression level *y_gi_(t)*, for gene *g* and cell *i* at time point *t* can be modeled as,

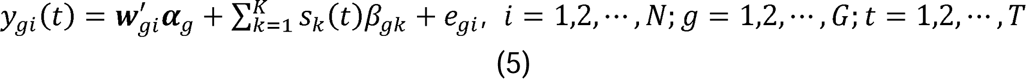

As a result, the parameter estimates of Equation 5 could fall into Equations 3 and 4, where the 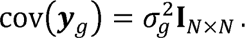. Therefore, we can directly apply the cone projection algorithm to estimate 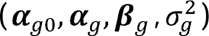 without iteration procedure. Compared with Mixed TDEseq, Linear TDEseq is much more efficient in analyzing large-scale time-course scRNA-seq data.

### Testing temporal expression patterns

Testing whether a gene shows a temporal expression pattern over all time points can be translated into testing the null hypothesis: ***H**_0_:**β**_g_* = **0**. The statistical Testing whether a gene shows a temporal expression pattern over all time points power of such a hypothesis test will depend on how well the pattern-constrained spline function fits the observed temporal expressions. We, therefore, compute *p*-values for growth, recession, peak, or trough (each at a time), thereby combining these four *p*-values through the Cauchy combination rule [45]. Specifically, we first convert each of the four *p*-values into a Cauchy statistic, and then aggregate the four Cauchy statistics through summation and convert the summation back to a single *p*-value based on the standard Cauchy distribution (Additional file 1: Supplementary Text). After obtaining *m p*-values across *m* genes, we computed the false discovery rate (FDR) through the permutation strategy.

### Determining one of the temporal expression patterns for each gene

A primary goal of TDEseq is assigning a suitable expression pattern of four given patterns (i.e., growth, recession, peak, or trough) for each gene. Specifically, for the linear version of TDEseq, TDEseq calculated the *Akaike* information criterion (AIC) [89] for gene *g*:

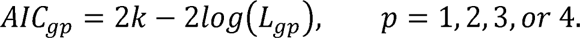

where *k* is the number of parameters and *L_gp_* is the likelihood for gene *g* and pattern *p*. For Mixed TDEseq, the marginal AIC is not an asymptotically unbiased estimator of the AIC and favors smaller models without random effects, but conditional AIC induces a bias that can lead to the selection of any random effect not predicted to be exactly zero [90]. Therefore, we used *B*-statistics to determine the temporal expression patterns for gene *g*:

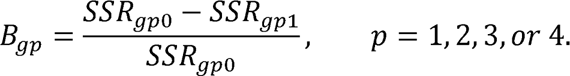

Where *SSR_gp1_* is the sum of squared residuals with parameters estimated via cone projection algorithm, and *SSR_gp0_* is the sum of squared residuals under the null hypothesis.

### Simulations

To make our simulations as realistic as possible, we simulated time-course scRNA-seq data using the *Splatter* package [46], in which the parameters were inferred from real LUAD scRNA-seq data [14]. *Splatter* simulated scRNA-seq data by specifying the number of cells (using *batchCells* parameter) and the number of time points (using *group.prob* parameter). Specifically, in null simulations, we set the probability of the differential expression genes *de.prob* = 0 and the effect of time point-specific size parameter *de.facloc* = 0 to denote the non-temporal expression gene across all stages. We simulated 200 cells measured by 10,000 genes for each sample, to examine the performance of type I error control.

In power simulations, we varied the number of time points, the number of cells per sample, the time point-specific effect size *de.facloc*, and expected UMI counts *lib.loc* (Additional file 3: Table S2). To do so, we set the probability of temporal expression gene in a group *de.prob* = 0.3, the number of stages as 5, the time point-specific effect size parameter *de.facloc* = 0.4, and the expected UMI count *lib.loc =* 9.4 as the baseline simulation scenario. We then varied the number of time points to be either 4, 5, or 6 (3 samples/replicates per time point); the number of cells to be either 100, 200, or 300 for each sample; the time point-specific effect size parameter as 0.1, 0.4 or 0.7; expected UMI counts as 7, 9.4 and 13.7 (estimates from rat liver scRNA-seq data). *Splatter* also added batch effects (i.e., sample-level variations) for the simulated data set. The batch effects were applied to all genes for each sample. For the batch effects, we simulated 100 cells for each batch, and we varied sample-level variations, i.e., batch effects (i.e., *batch.facloc*) to be either 0, 0.04, or 0.12 to represent small, moderate, or strong batch effects. With these parameter settings, we limited our simulations to six specific temporal expression patterns ---- growth, recession, peak, trough, bi-plateau, and multi-modal patterns. For temporal expression effect sizes, we generated time point-specific effect sizes by setting the parameter *de.facloc*, one time point at a time. Then, we examined temporal expression patterns based on effect sizes across time points, limiting our simulations to six specific temporal expression patterns (Additional file 2: Fig. S1). We simulated three samples/replicates for each time point and repeated 10 times for each simulation scenario.

Besides temporal expression patterns, we also generated the smudged time-course scRNA-seq data using the *SymSim* R package [58]. Specifically, the parameter settings were as follows: the transcription rate *vary* = “s”; the variance of Brownian motion, *Sigma* = 0.4; the mean rate of subsampling of transcripts *alpha_mean* = 0.05; the standard deviation rate of subsampling of transcripts *alpha_sd* = 0.02; the mean of sequencing depth *depth_mean* = 10,000; and the standard deviation of sequencing depth *depth_sd* = 3,000. All data sets were measured by 2,000 cells and 10,000 genes. Consequently, to generate the data that were intertwined cells among time points, we proportionally mixed the cells from the other two adjacent time points for each stage. Specifically, with a given time point, we randomly sampled *p*_1_ = 90% cells from the given stage, then sampled *p*_2_ 8% and *p*_3_ = 2% from two adjacent stages, respectively, as a low proportion of intertwined cell scenario. Similarly, we set *p*_1_ = 70%, *p*_2_ = 24% and *p*_3_ = 6% as a medium proportion of intertwined cell scenario and *p*_1_ = 50% *p*_2_ 40% and *p*_3_ = 10% as a high proportion of intertwined cell scenario.

### The difference in statistical models among TDEseq and tradeSeq or ImpluseDE2

In addition to TDEseq, we also employed other two temporal expression analysis methods, tradeSeq, and ImpulseDE2. The tradeSeq builds on the generalized additive model (GAM) that directly models raw gene expression counts in scRNA-seq data, i.e.,

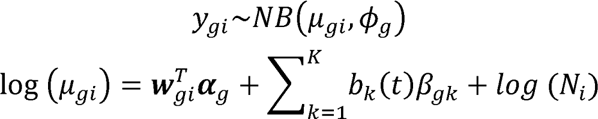

where *y_gi_* represents the raw counts for gene *g* and cell *i*; *t* represents the time points; ***w**_gi_* represents the covariates and 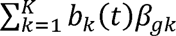 represents a linear combination of *K* cubic basis functions; *N_i_* denotes the total counts for cell *i; NB* is a negative binomial distribution. To avoid overfitting issues, tradeSeq employed penalized spline which shrinkages *β_gk_* to zero, and therefore are less sensitive to temporal genes with small fold changes. Notably, tradeSeq [47] was primarily developed for detecting trajectory-based differential expression genes, however, the applicability of tradeSeq was extended beyond this setting, i.e., also can be applied to bulk time-course RNA-seq data analysis [31]. Therefore, we performed an analogous analysis using tradeSeq in the comparison. On the other hand, ImpulseDE2 [25] combines the impulse model [82] with a negative binomial noise model to directly model the raw counts of gene expression measurements. The impulse function is the scaled product of two sigmoid functions:

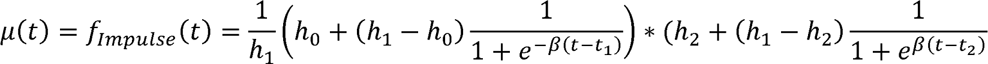

where 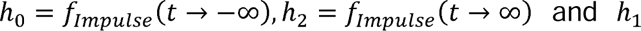 models the intermediate expression, t1 and t2 are the state transition times, *β* is the slope parameter of both sigmoid functions.

The impulse function is a more restrictive model compared with spline functions, therefore limiting its power. It was originally designed to model the bulk time-course RNA-seq data. To adapt for temporal expression analysis of time-course scRNA-seq data, we modified the implementation of ImpulseDE2 following the tradeSeq paper. Both methods take a count matrix ***Y*** and a time points vector ***t*** as input and return one *p*-value for each gene at a time.

## Methods for comparison

We compared TDEseq with five existing methods for identifying temporal expression genes from time-course from scRNA-seq data (tradeSeq, and Wilcoxon test) or bulk RNA-seq data (ImpulseDE2, edgeR, and DESeq2). For tradeSeq (version 1.4.0), we used the functions *fitGAM* and *associationTest* (https://statomics.github.io/tradeSeq/articles/tradeSeq.html). The number of knots parameter *k* in the tradeSeq was chosen by 100 random genes based on the tradeSeq vignette. For ImpulseDE2 (version 0.99.10), we followed the modified implementation of ImpulseDE2 in the tradeSeq paper (https://github.com/statOmics/tradeSeqPaper). For DESeq2 (version 1.40.2) and edgeR (version 3.42.4), we treated time points as categorical factors and tested DE genes using a likelihood ratio test.

### Permutation strategy to construct the null distribution

In real data applications, to calculate the false discovery rate (FDR), we construct an empirical null distribution of *p*-values through permuting the time point variables and repeating 5 times. Afterward, we computed the FDRs using

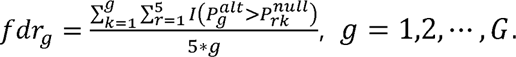

where 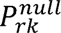 is an increasing ordered *p*-value for, *k^th^* gene and *k^th^* permutation; 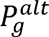 is an increasing ordered *p*-values under the alternative hypothesis.

### Functional gene set enrichment analysis

The gene set enrichment analyses of temporal expression genes were performed by the *enrichGO* function implemented in the *clusterProfiler* R package (version 3.18.1) [91]. Specifically, we used all genes as the background and set the minimal and maximal sizes of genes annotated by Gene Ontology (GO) terms for testing as 10 and 500, respectively. The significant GO terms were selected by setting BH-adjusted *p*-value < 0.05.

### Batch effects removal evaluation

We used the LISI metric to measure cell batch distribution (iLISI) [57]. The LISI metric was designed to assess whether clusters of cells in a scRNA-seq dataset are well-mixed across categorical variables (batches). We took the median value of the scores computed for all cells in the dataset and scaled the value between 0 and 1 to denote the worst and best cell mixed.

### HCT116 cell lines after 5-AZA-CdR treatment data

The scRNA-seq data were assayed by Well-TEMP-seq, which contains 5-AZA-CdR-treatment HCT116 cell lines after 0 days (4,000 cells), 1 day (4,000 cells), 2 days (4,000 cells) or 3 days (4,000 cells) [23]. The Well-TEMP-seq technology can distinguish new RNAs from pre-existing RNAs, we used the new RNAs which better reflect RNA dynamics for downstream analysis. For the preprocessing of scRNA-seq data, the genes with more than 99% zero counts were filtered out, resulting in 7,314 genes and 16,000 cells for further analysis.

### Mouse hepatoblast differentiation data

The scRNA-seq data were assayed by Smart-seq2 protocols [64] on isolated cells from mouse fetal livers at 7 different developmental stages [38]. Gene expression levels were measured by a total of 14,226 genes and 345 cells. In our analysis, we only considered the hepatoblast cells for temporal expression analysis. Finally, for the preprocessing of scRNA-seq data, the genes with more than 99% of zero counts were filtered out, resulting in 14,180 genes and 345 cells for further analysis.

### Human metastatic LUAD progression data

The scRNA-seq data [14] were assayed by 10X Genomics Chromium protocols [92] on LUAD samples from 5 distinct developmental stages, i.e., the control stage consists of a total of 80,441 cells in 7 cell types from 21 samples; stage I consists of a total of 31,026 cells in 7 cell types from 8 samples; stage II consists of a total of 3,840 cells in 7 cell types from 1 sample, stage III consists of a total of 10,283 cells in 7 cell types from 2 samples, and stage IV (metastasis) consists of a total of 82,916 cells in 7 cell types from 26 samples. In our analysis, we only employed the epithelial cells from the control lung samples (3,703 cells), stage I tumor lung samples (5,651 cells), stage III tumor lung samples (1,500 cells), and stage IV samples (lymph node metastasis, 6,582 cells). To relieve the computational burden in practice, we utilized a down-sampling strategy to randomly select 1,000 cells for the stage that contains more than 1,000 cells. Finally, for the preprocessing of scRNA-seq data, the genes with more than 99% of zero counts were filtered out, resulting in 15,263 genes and 4,000 cells for further analysis.

### Human COVID-19 immune response data

The scRNA-seq data were assayed by 10X Genomics Chromium protocols on human SARS-COV-2 infection samples from disease progression ranging from 4 days to 123 days [9]. In our analysis, we only employed the NK cells from the 21 serve/critical patients and divided those patients into 5 stages according to the time point interval, i.e. stage I (4-8 days, 930 cells), stage II (10-13days, 939 cells), stage III (19-24 days, 893 cells), stage IV (28-34days, 768 cells) and stage V (110-123days, 1,000 cells). Finally, for the preprocessing of scRNA-seq data, the genes with more than 99% of zero counts were filtered out, resulting in 10,699 genes and 4,530 cells for further analysis.

## AVAILABILITY OF DATA AND CODE

All scRNA-seq data sets used in this study are publicly available on the GEO database. Specifically, mouse hepatoblast differentiation scRNA-seq data are available at GEO under accession GSE90047; Human metastatic LUAD progression scRNA-seq data are available at GEO under accession GSE131907; Human COVID-19 immune response scRNA-seq data are available at GEO under accession GSE158055. Well-TEMP-seq scRNA-seq data are available at GEO under accession GSE194357. TDEseq is an open-source R package that is freely available from GitHub[93] (https://github.com/fanyue322/TDEseq) and https://sqsun.github.io/software.html. Source code for the software release used in the paper has been placed into a DOI-assigning repository[94](https://zenodo.org/records/10869078). The source code and scRNA-seq data for reproducing the results are publicly available at https://sqsun.github.io/software.html.

## ACKNOWLEDGEMENTS

We would like to thank Jing Ning at Xi’an Jiaotong University for her help processing part of the data used in this paper.

## CONTRIBUTIONS

SS conceived the idea of the manuscript and provided funding support. SS and YF developed the method and designed the experiments. YF implemented the software and performed simulations and real data analysis with assistance from

LL. SS and YF wrote the manuscript. All authors approved the final manuscript.

## Ethics declarations

Not applicable

---

**Additional file 1:**

Supplementary text on TDEseq modeling and inference details.

**Additional file 2:**

Supplementary figures on the simulation performance evaluation.

**Additional file 3:**

Supplementary tables on simulations and real data information, identified dynamic temporal genes for each real dataset. Additionally, the validation gene sets utilized in this research are documented.

