## Additional file 1 for "Powerful and accurate detection of temporal gene expression patterns from multi-sample multi-stage single cell transcriptomics data with TDEseq"

**Supplementary Text**

**Inference algorithm of TDEseq**

Here, we provide details on the parameter estimation of TDEseq models. All notations follow the Overview of TDEseq section in the main text. Specifically, we recall that

$\mathbf{E}\left( \boldsymbol{y}_{g} \right)=\mathbf{W}\boldsymbol{\alpha}_{g}+\left[ \mathbf{W}_{0}\mathbf{S} \right]\mathbf{[}\begin{matrix} \boldsymbol{\alpha}_{g0} \\ \boldsymbol{\beta}_{g} \end{matrix}\mathbf{]}\boldsymbol{.}$ $g=1,2,\cdots,G$.

(1)

where $\mathbf{S}$is an $N\times m$ matrix with columns $s_{1}\left( t \right),\ldots,s_{k}\left( t \right)$, and $k$ is the number of knots of the spline function. $\mathbf{W}$ is the covariates matrix; $\boldsymbol{\alpha}_{g}$**,** $\boldsymbol{\alpha}_{g0}$ and$\boldsymbol{\beta}_{g}$are the corresponding regression coefficients. For the growth or recession patterns,$\mathbf{S}$ was assigned by *I*-spline basis functions and $\mathbf{W}_{0}=\left[ \mathbf{1}_{N} \right]$. For both peak and trough patterns, $\mathbf{S}$ was assigned by the *C*-spline basis vectors and $\mathbf{W}_{0}=[\mathbf{1}_{N},\boldsymbol{t]}$, where $\boldsymbol{t}=\left( t_{1},\ldots,t_{N} \right)^{'}$ represents the time points.

For any pattern constraints, we assume a linearly independent set of $\mathbf{S,}\mathbf{W}_{0}$and $\mathbf{W}$ together as a closed convex cone:

$$\nabla=\left\{ \boldsymbol{\mu}_{g}\in\mathbb{R}^{N}:\boldsymbol{\mu}_{g}\boldsymbol{=}\mathbf{E}\left( \boldsymbol{y}_{g} \right)=\mathbf{W}_{0}\boldsymbol{\alpha}_{g0}+\mathbf{W}\boldsymbol{\alpha}_{g}+\mathbf{S}\boldsymbol{\beta}_{g},\boldsymbol{\beta}_{g}\geq0 \right\}$$

(2)

The least-squares estimator for $\boldsymbol{\mu}_{g}$ is the projection of $\boldsymbol{y}_{g}$ onto $\nabla$. For linear TDEseq, note that $\mathrm{cov}\left( \boldsymbol{y}_{\boldsymbol{g}} \right)=\sigma^{2}\mathbf{I}$, we estimate parameters ${\hat{\boldsymbol{\alpha}}}_{\boldsymbol{g}\boldsymbol{0}}, {\hat{\boldsymbol{\alpha}}}_{\boldsymbol{g}},$ ${\hat{\boldsymbol{\beta}}}_{g}$ and $\hat{\sigma}^{2}$ through cone projection algorithm relies on the *coneproj* package in R [1]. For Mixed TDEseq, instead of the $\boldsymbol{y}_{\boldsymbol{g}}$ being IID, we assume gene expression levels are correlated due to batch effects, i.e., $\mathrm{cov}\left( \boldsymbol{y}_{\boldsymbol{g}} \right)={\sigma_{g}^{2}\mathbf{K}}_{\boldsymbol{g}}$, $\mathbf{K}_{g}\boldsymbol{=}\frac{\sigma_{gu}^{2}}{\sigma_{g}^{2}}\boldsymbol{\Sigma}_{N\times N}+\mathbf{I}_{N\times N}$, while cone projection algorithm requires uncorrelated $\boldsymbol{y}_{\boldsymbol{g}}$. If the positive definite matrix $\mathbf{K}_{\boldsymbol{g}}$ is known and $\mathbf{K}_{\boldsymbol{g}}\boldsymbol{=}\mathbf{P}_{\boldsymbol{g}}\boldsymbol{\Lambda}_{\boldsymbol{g}}\mathbf{P}_{\boldsymbol{g}}=\left( \mathbf{P}_{\boldsymbol{g}}\sqrt{\boldsymbol{\Lambda}_{\boldsymbol{g}}} \right)\left( \sqrt{\boldsymbol{\Lambda}_{\boldsymbol{g}}}\mathbf{P}_{\boldsymbol{g}}^{'} \right)=\mathbf{U}_{\boldsymbol{g}}\mathbf{U}_{\boldsymbol{g}}^{'}$ is the Eigen decomposition of $\mathbf{K}_{\boldsymbol{g}}$, then we transform the variable of Equation 2 as: ${\tilde{\boldsymbol{y}}}_{g}\mathbf{=}\mathbf{U}_{g}^{\boldsymbol{-1}}\boldsymbol{y}_{g}\mathbf{,}{\tilde{\mathbf{W}}}_{0}\mathbf{=}\mathbf{U}_{g}^{\boldsymbol{-1}}\mathbf{W}_{0}\mathbf{,}\tilde{\mathbf{W}}\mathbf{=}\mathbf{U}_{g}^{\boldsymbol{-1}}\mathbf{W}$, and$\mathbf{,} \tilde{\boldsymbol{S}}\mathbf{=}\mathbf{U}_{g}^{\boldsymbol{-1}}\mathbf{S}$, resulting in

$$\nabla=\left\{ {\tilde{\boldsymbol{\mu}}}_{g}\in\mathbb{R}^{N}:{\tilde{\boldsymbol{\mu}}}_{g}={\tilde{\mathbf{W}}}_{0}\boldsymbol{\alpha}_{g0}+\tilde{\mathbf{W}}\boldsymbol{\alpha}_{g}+\tilde{\mathbf{S}}\boldsymbol{\beta}_{g},\boldsymbol{\beta}_{g}\geq0 \right\}$$

(3)

Because $\mathrm{cov}\left( {\tilde{\boldsymbol{y}}}_{g} \right)$=$\sigma_{g}^{2}\mathbf{I}$, we can perform the cone projection and infer the parameters on the log-normalized gene expression levels of scRNA-seq data.

For simplicity, we dropped the subscript $g$ in the following notations. Here, we denote $\theta=\frac{\sigma_{u}^{2}}{\sigma^{2}}$ and $\boldsymbol{y}=\left( y_{1,1},\ldots,y_{1,n_{1}},\ldots,y_{M,1},\ldots,y_{M,n_{M}} \right)^{'}$, and $n_{j}$is the number of cells for each sample; $j=1,\ldots,M$ represents the index of individuals, $\mathbf{K}$ is a block diagonal matrix that the $jth$ block of $\mathbf{K}$ is:

$$\mathbf{K}_{\boldsymbol{j}}\left( \theta\right)=\left( \begin{matrix} \begin{matrix} \theta+1 & \theta\\ \theta& \theta+1 \end{matrix} & \cdots& \begin{matrix} \theta\\ \theta\end{matrix} \\ \vdots& \ddots& \vdots\\ \begin{matrix} \theta& \theta\end{matrix} & \cdots& \theta+1 \end{matrix} \right)$$

and $\mathbf{K}\boldsymbol{=}diag\{\mathbf{K}_{\boldsymbol{1}}\left( \theta\right),\ldots,\mathbf{K}_{\boldsymbol{M}}\left( \theta\right)\}$.

To estimate $\theta$, we write the log-restricted likelihood of $\sigma^{2}$ and $\theta$:

$$l_{r}\left( \sigma^{2},\theta\right)\propto-\frac{1}{2}log|\sigma^{-2}\mathbf{X}^{'}\mathbf{K}^{-1}\left( \theta\right)\mathbf{X}|-\frac{1}{2}\sigma^{-2}\sum_{j=1}^{N} \left( y_{ij}-\mu_{ij} \right)^{'}\mathbf{K}^{\mathbf{-1}}\left( \theta\right)\left( y_{ij}-\mu_{ij} \right)-\frac{1}{2}log|\sigma^{2}\mathbf{K}\left( \theta\right)|$$

where $\mathbf{X}=[\mathbf{W}_{0},\mathbf{W}]$. Assume $\mu_{ij}$ is known, we can approximate $\sigma^{2}$ as:

$$\hat{\sigma}^{2}\left( \theta\right)\approx\frac{1}{N-d}\left( \boldsymbol{y}-\hat{\boldsymbol{\mu}} \right)^{'}\mathbf{K}^{-1}\left( \theta\right)\left( \boldsymbol{y}-\hat{\boldsymbol{\mu}} \right)$$

(4)

Where $d$ is a measure of model degrees of freedom that are also estimated through the cone projection algorithm. Given $\mu_{ji}$ and $\sigma^{2}$, after performing some algebraic operations, we can estimate $\hat{\theta}$ by calculating the root of the first derivation of $L\left( \theta\right)$:

$$L^{'}\left( \theta\right)=\frac{N-P}{2}\frac{\sum_{j=1}^{M} \frac{\left( n_{j}\bar{r}_{j} \right)^{2}}{\left( 1+n_{j}\theta\right)^{2}}}{\left| \left| \boldsymbol{r} \right| \right|^{2}-\sum_{j=1}^{M} \frac{\theta\left( n_{j}\bar{r}_{j} \right)^{2}}{1+n_{j}\theta}}-\frac{1}{2}\sum_{j=1}^{M} \frac{n_{j}}{1+n_{j}\theta}+tr(\left( \mathbf{X}^{'}\mathbf{K}\left( \theta\right)\mathbf{X} \right)^{-1}\mathbf{X}^{'}\mathbf{1}_{N\times N}\mathbf{X})$$

(5)

Where $r_{ji}=y_{ji}-\mu_{ji}$ is the residuals and $\bar{r}_{j}$ is the average residual for all cells belonging to individual $j$, $\mathbf{1}_{N\times N}$ is an $N\times N$ all-ones matrix, and $N=\sum_{j=1}^{M} n_{j}$ is the total number of cells.

The designed LAMM inference algorithm is implemented by iteratively estimating ${\hat{\boldsymbol{\alpha}}}_{\boldsymbol{0}}, \hat{\boldsymbol{\alpha}},$ $\hat{\boldsymbol{\beta}}$, $\hat{\sigma}^{2}$ and $\hat{\theta}$ until convergence is reached, utilizing equations (3), (4) and (5), the algorithm is summarized in **Algorithm 1**:

**Algorithm 1**. The parameter inference algorithm for Mixed TDEseq

**Input:** Normalized gene expression data $\boldsymbol{y}=\left( y_{1,1},\ldots,y_{1,n_{1}},\ldots,y_{M,1},\ldots,y_{M,n_{M}} \right)^{'}$, the spline basis function $\left[ \mathbf{W}_{0}\mathbf{S} \right]$ and covariates $\mathbf{W}$, maximum iterations $maxIter$, relative tolerance $tol$.

**Output:** ${\hat{\boldsymbol{\alpha}}}_{\boldsymbol{0}}, \hat{\boldsymbol{\alpha}},$ $\hat{\boldsymbol{\beta}}$, $\hat{\sigma}^{2}$ and $\hat{\theta}$

1: Initialize $\hat{\theta}^{(0)}=0$, estimate ${\hat{\boldsymbol{\alpha}}}_{\boldsymbol{0}}^{\boldsymbol{(0)}}, {\hat{\boldsymbol{\alpha}}}^{\boldsymbol{(0)}}, {\hat{\boldsymbol{\beta}}}^{(0)},$based on Equation 3 using a cone projection algorithm.
2: **for** $it\in1,\ldots,maxIter$ **do**
3: Update model parameters $\hat{\sigma}_{(it)}^{2}$ based on Equation 4.
4: Update model parameters $\hat{\theta}^{(it)}$ based on Equation 5.
5: Perform eigen decomposition $\mathbf{K}^{\boldsymbol{(}it\boldsymbol{)}}=\mathbf{U}\mathbf{U}^{T}$, transformed $\tilde{\boldsymbol{y}}\mathbf{=}\mathbf{U}^{\boldsymbol{-1}}\boldsymbol{y}\mathbf{,}{\tilde{\mathbf{W}}}_{0}\mathbf{=}\mathbf{U}^{\boldsymbol{-1}}\mathbf{W}_{0}\mathbf{,}\tilde{\mathbf{W}}\mathbf{=}\mathbf{U}^{\boldsymbol{-1}}\mathbf{W}$, and $\tilde{\mathbf{S}}\mathbf{=}\mathbf{U}^{\boldsymbol{-1}}\mathbf{S}$.
6: Update model parameters ${\hat{\boldsymbol{\alpha}}}_{\boldsymbol{0}}^{\boldsymbol{(}it\boldsymbol{)}}, {\hat{\boldsymbol{\alpha}}}^{\boldsymbol{(}it\boldsymbol{)}}, {\hat{\boldsymbol{\beta}}}^{(it)}$ based on Equation 3 using the cone projection algorithm.
7: Evaluate $\boldsymbol{\mu}^{\left( it \right)}\boldsymbol{=}{\tilde{\mathbf{W}}}_{0}{\hat{\boldsymbol{\alpha}}}_{\boldsymbol{0}}^{\left( it \right)}+\tilde{\mathbf{W}}{\hat{\boldsymbol{\alpha}}}^{\left( it \right)}+\tilde{\mathbf{S}}{\hat{\boldsymbol{\beta}}}^{\left( it \right)}$
8: **if** $\left| \boldsymbol{\mu}^{\left( it \right)}\boldsymbol{-}\boldsymbol{\mu}^{\left( it\boldsymbol{-1} \right)} \right|\boldsymbol{<}tol$
9: break;
10: **end if**
11: **end for**
12: **return** ${\hat{\boldsymbol{\alpha}}}_{\boldsymbol{0}}, \hat{\boldsymbol{\alpha}},$ $\hat{\boldsymbol{\beta}}$, $\hat{\sigma}^{2}$ and $\hat{\theta}$

**Statistical testing and *p*-value calculation**

To test if the gene $g$ is a temporal expression gene over time points, TDEseq defines the null and alternative hypotheses as:

$$\mathbf{H}_{0}: \boldsymbol{\beta}_{g}=\boldsymbol{0}, \mathbf{H}_{1}: \boldsymbol{\beta}_{g}\geq\boldsymbol{0}$$

denote $SSR_{p0}$ is the sum of squared residuals under $\mathbf{H}_{0}$, and $SSR_{p1}$ is the sum of squared residuals under $\mathbf{H}_{1}$, then

$$B_{p}=\frac{SSR_{p0}-SSR_{p1}}{SSR_{p0}}, p=1, 2, 3, or 4.$$

The large values of $B_{p}$ support the alternative hypothesis. A standard result in cone projection is that the null distribution of $B_{p}$ is a mixture of beta random variables [2]. Specifically, when $\mathbf{H}_{0}$ is true, then $B_{p}$ follow the distribution as follows:

$$\Pr\left( B_{p}\leq a \right)=\sum_{d=0}^{k} \Pr\left[ B\left( \frac{d}{2},\frac{N-d}{2} \right)\leq a \right]p_{d}$$

where $B\left( \frac{d}{2},\frac{N-d}{2} \right)$is a random variable generated from $Beta\left( \frac{d}{2},\frac{N-d}{2} \right)$, and $k$ is the number of knots of the spline function. To obtain $p_{d}$, we randomly simulated $\check{y}\sim MVN(0,\boldsymbol{I}_{N})$ and performed cone projection algorithm with $\check{\boldsymbol{y}}$ and $({\tilde{\mathbf{W}}}_{0}, \tilde{\mathbf{W}}\boldsymbol{,} \tilde{\mathbf{S}}\boldsymbol{)}$ obtained from the last iteration in **Algorithm 1**. We then performed a cone projection algorithm for each $\check{\boldsymbol{y}}$ to obtain an empirical distribution of $p_{d}$. We then calculated the *p*-values according to such mixture *Beta* distribution.

After obtaining 4 different *p*-values, one for each pattern (i.e., growth, recession, peak, and trough), we combined these *p*-values into a single *p*-values through the Cauchy combination rule [3]. Specifically, we converted each of the 4 *p*-values into Cauchy statistics, averaged the four Cauchy statistics, and then converted the average back to a single *p*-value using the Cauchy distribution. The Cauchy rule takes advantage of the fact that the summation of Cauchy random variables also follows a Cauchy distribution regardless of whether these random variables are correlated or not. Therefore, the Cauchy rule allows us to combine multiple potentially correlated *p*-values into a single *p*-value without loss of type I error control. Due to the numerical precision of the Cauchy cumulative density function, the combined *p*-value is set as zero when it is below the threshold of 5.55e-17. Finally, after obtaining *G* *p*-values across *G* genes, we adjusted the *p*-value using FDR correction.
