## Additional file 2 for "Powerful and accurate detection of temporal gene expression patterns from multi-sample multi-stage single cell transcriptomics data with TDEseq"

**Supplementary Figures**

**Fig. S1. The temporal expression patterns in simulations.** The temporal expression patterns were simulated using *Splatter*. We generated time point-specific effect sizes by setting the parameter *de.facloc*, one time point at a time. Consequently, we examined temporal expression patterns based on effect sizes across time points, limiting our simulations to six specific temporal expression patterns, i.e., (**A**) growth pattern; (**B**) recession pattern; (**C**) peak pattern; (**D**) trough pattern; (**E**) bi-plateau pattern; (**F**) multi-modal pattern. The different colors denote the different time points.

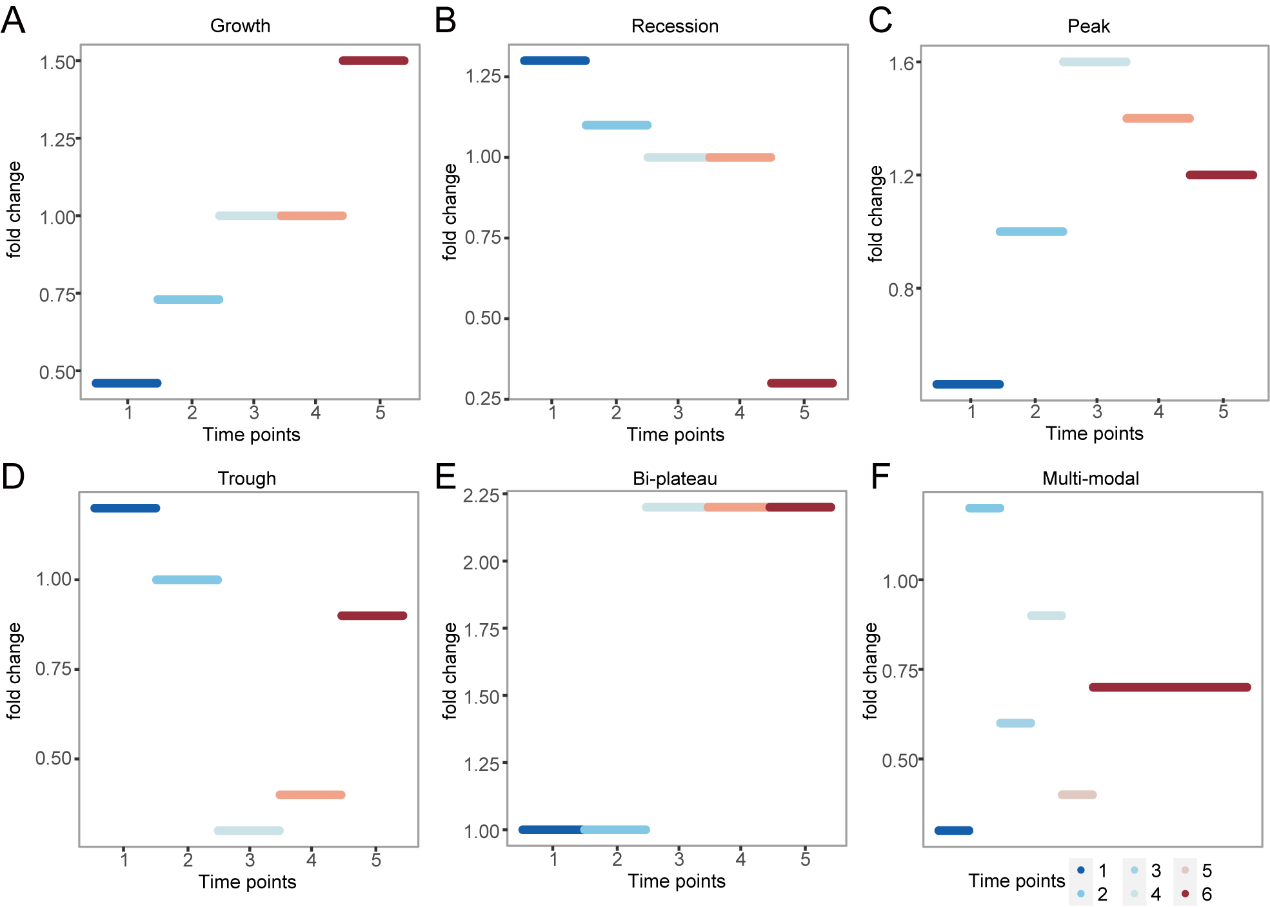

**Fig. S2.** **The performance of temporal gene detection methods is evaluated via varying the number of time points in simulations.** The quantile-quantile (QQ) plot shows the type I error control when the number of time points is equal to 4 (**A**) or 6 (**B**). The well-calibrated *p*-values will be expected laid on the diagonal line. Mixed TDEseq generated the well-calibrated *p-*values in both scenarios. generated from produced the *p*-values that are not well calibrated. The power comparison of temporal expression gene detection across a range of FDR cutoffs over 4 time points (**C**) or 6 time points (**D**). Both versions of TDEseq exhibit high temporal expression gene detection power, followed by DESeq2, edgeR, tradeSeq, and ImpulseDE2. FDR: false discovery rate. We benchmark a total of seven methods, i.e., Mixed TDEseq (plum), Linear TDEseq (orange) and DESeq2 (brown), tradeSeq (green), ImpulseDE2 (blue), edgeR (dark green), and Wilcoxon test (yellow). The TDEseq methods were highlighted using solid lines, while other methods were represented by dashed lines in the plots.

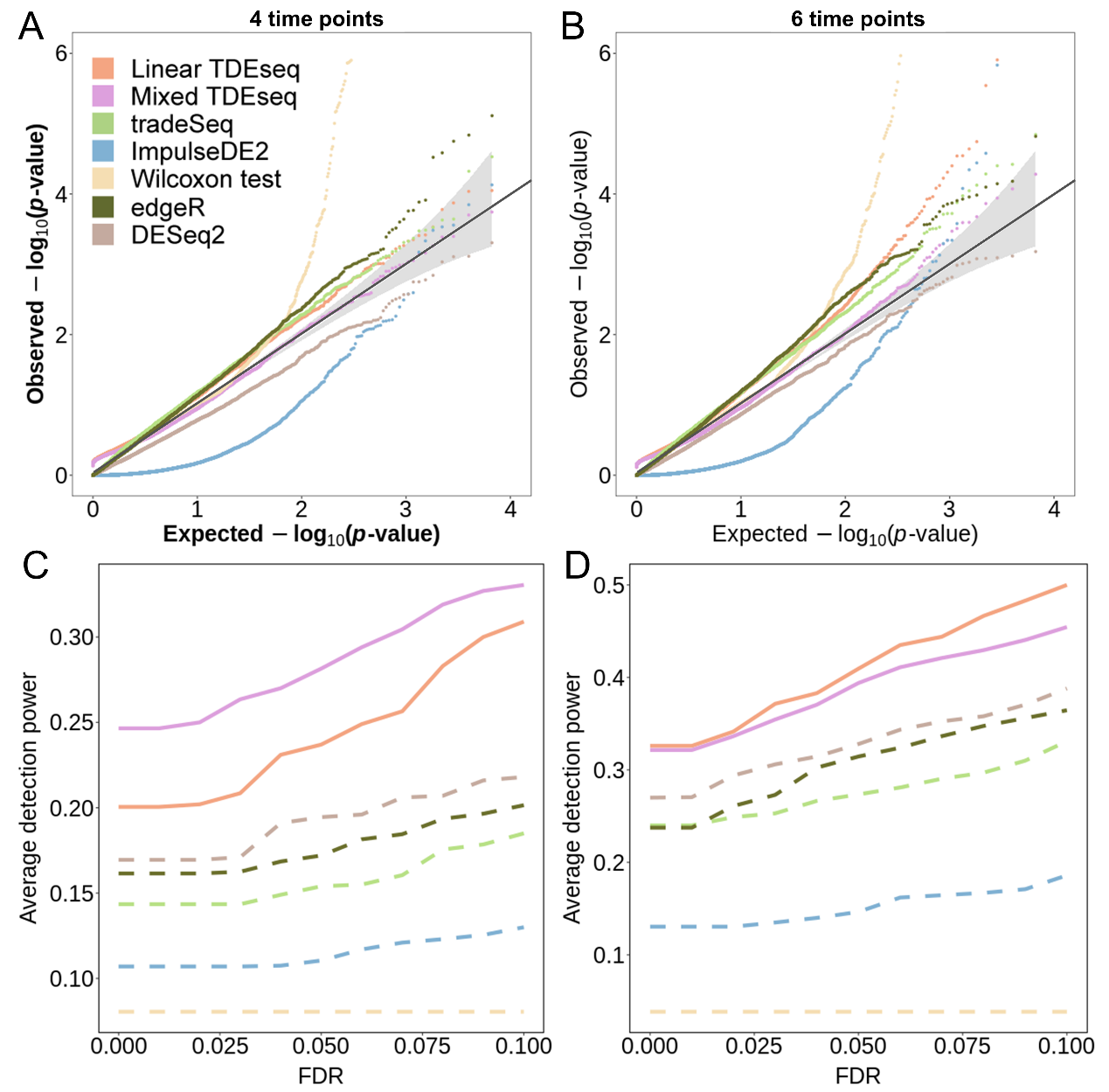

**Fig. S3.** **The performance of temporal gene detection methods is evaluated via varying the number of expected UMI counts for each cell in simulations.** The quantile-quantile (QQ) plot shows the type I error control with (**A**) low UMI counts (*lib.loc =* 7.0) and (**B**) high UMI counts (*lib.loc =* 13.8). TDEseq generated well-calibrated *p*-values in low UMI counts but not in high UMI counts. The power comparison of temporal expression gene detection across a range of FDR cutoffs with (**C**) low UMI counts or (**D**) high UMI counts. Both versions of TDEseq exhibit high temporal expression gene detection power under the low UMI counts scenario, while pseudo-bulk-based methods exhibit high temporal expression gene detection power under the high UMI counts scenario. We benchmark a total of seven methods, i.e., Mixed TDEseq (plum), Linear TDEseq (orange) and DESeq2 (brown), tradeSeq (green), ImpulseDE2 (blue), edgeR (dark green), and Wilcoxon test (yellow). The TDEseq methods were highlighted using solid lines, while other methods were represented by dashed lines in the plots.

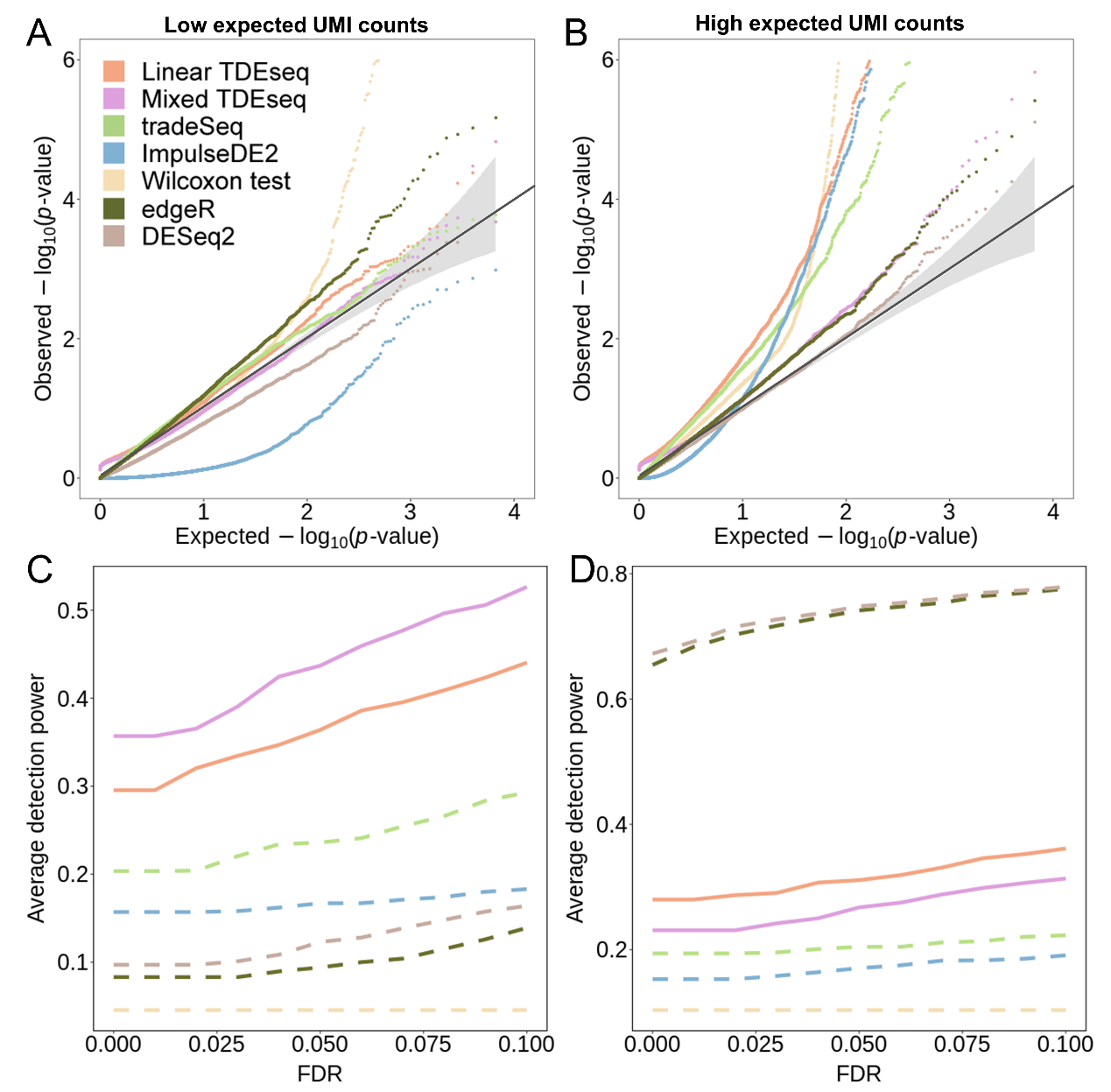

**Fig. S4.** **The performance of temporal gene detection methods is evaluated via varying the number of cells per sample in simulations.** The quantile-quantile (QQ) plot shows the type I error control with (**A**) a small number of cells (100 cells per sample) and (**B**) a large number of cells (300 cells per sample). TDEseq generated the reasonably well-calibrated *p*-values in both scenarios. The power comparison of temporal expression gene detection across a range of FDR cutoffs with a small number of cells per sample (**C**) and a large number of cells per sample (**D**). Both versions of TDEseq exhibit high temporal expression gene detection power under a large number of cells per sample scenario, while pseudo-bulk-based methods exhibit high temporal expression gene detection power under a small number of cells per sample scenario. We benchmark a total of seven methods, i.e., Mixed TDEseq (plum), Linear TDEseq (orange) and DESeq2 (brown), tradeSeq (green), ImpulseDE2 (blue), edgeR (dark green), and Wilcoxon test (yellow). The TDEseq methods were highlighted using solid lines, while other methods were represented by dashed lines in the plots.

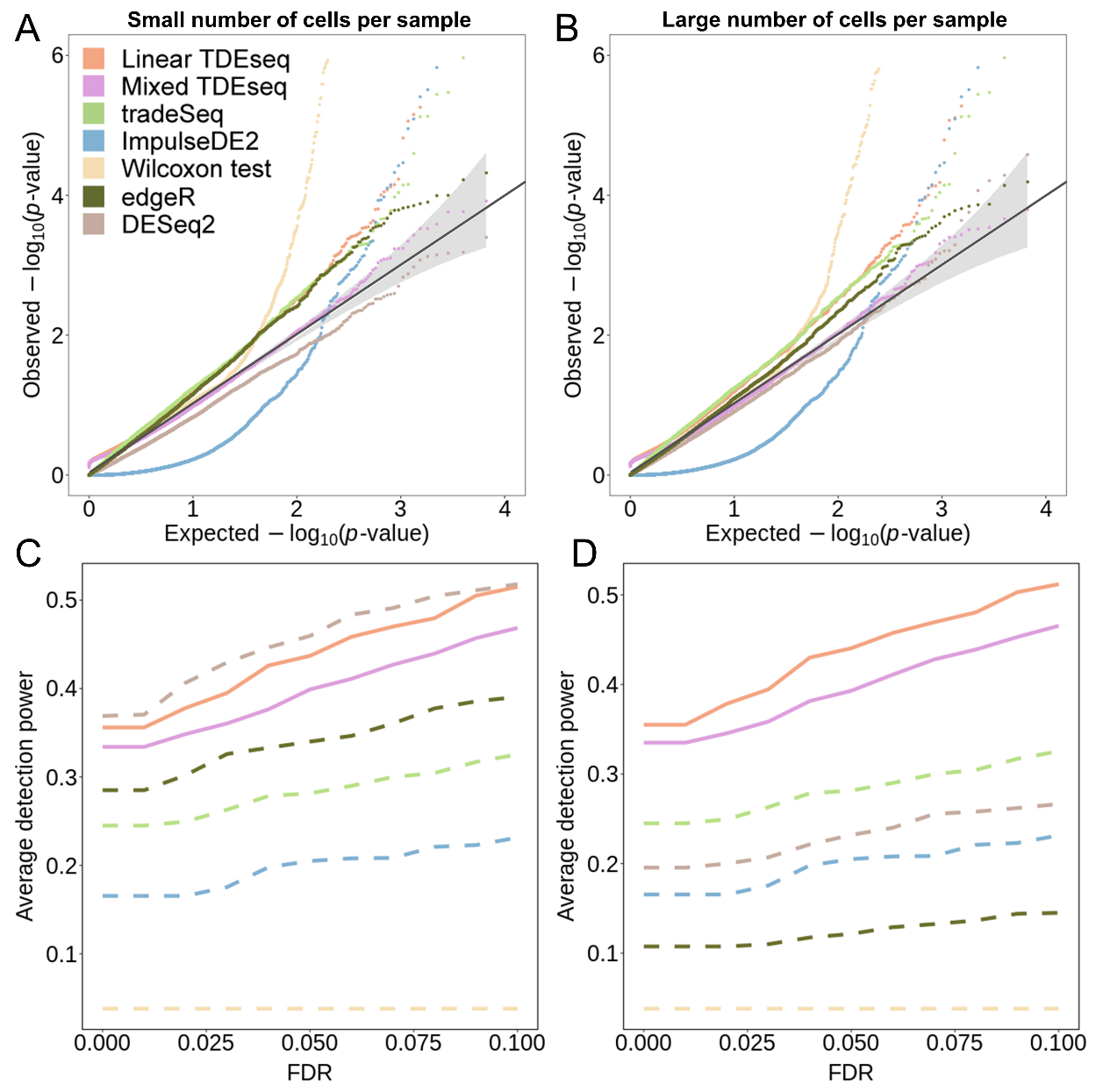

**Fig. S5.** **The performance of temporal gene detection methods is evaluated via varying the effect size in simulations.** The power comparison of temporal expression gene detection across a range of FDR cutoffs under (**A**) small effect size (*de.facloc* = 0.1) and (**B**) large effect size (*de.facloc* = 0.7). TDEseq exhibits high temporal expression gene detection power. We benchmark a total of seven methods, i.e., Mixed TDEseq (plum), Linear TDEseq (orange) and DESeq2 (brown), tradeSeq (green), ImpulseDE2 (blue), edgeR (dark green), and Wilcoxon test (yellow). The TDEseq methods were highlighted using solid lines, while other methods were represented by dashed lines in the plots. FDR: false discovery rate.

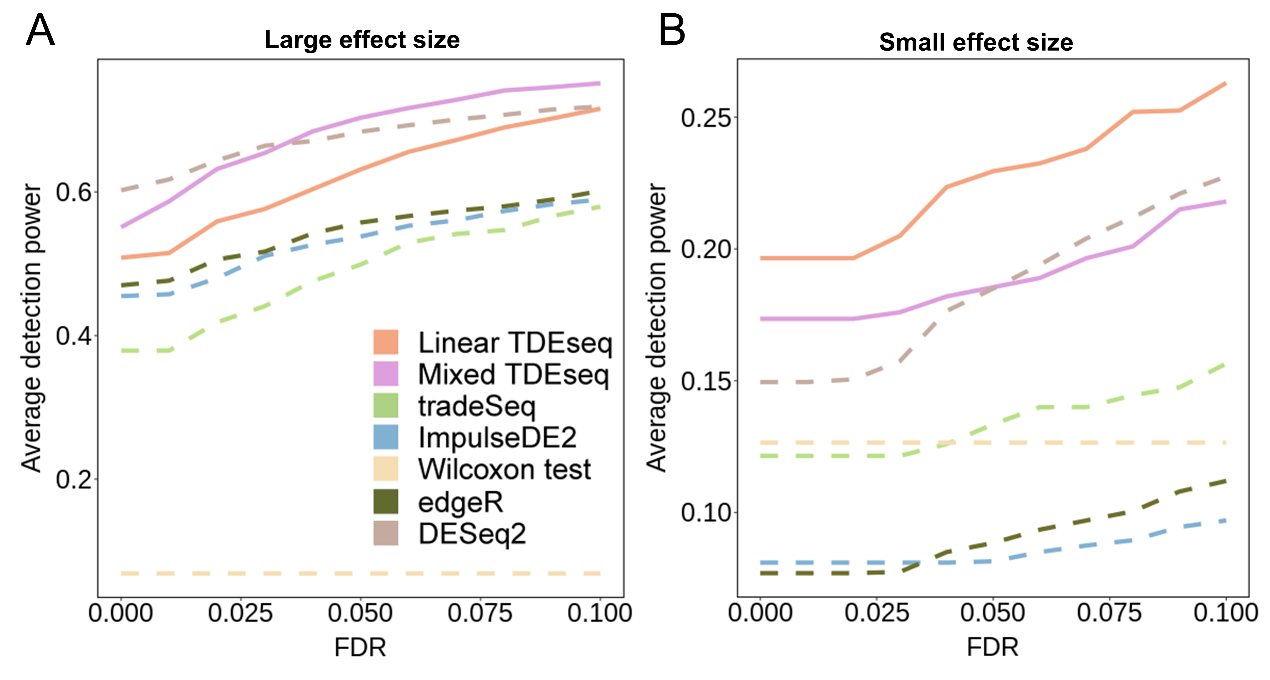

**Fig. S6.** **The comparison of the accuracy of pattern-specific detection with varying different parameter settings in simulations.** (**A**) When the number of time points is 4. (**B**) When the number of time points is 6. (**C**) When the effect size is small (0.1). (**D**) When the effect size is large (0.7). (**E**) When the expected UMI counts per cell are low (7.0). (**F**) When the expected UMI counts per cell are high (13.8). (**G**) When the number of cells per sample is small (50 cells in each sample). (**H**) When the number of cells per sample is large (200 cells in each sample). (**I**) When there are no batch effects. We found Mixed TDEseq (plum) outperforms ImpulseDE2 (blue) in all parameter settings. The tradeSeq, Wilcoxon test DESeq2, and edgeR are excluded from this comparison due to there being no pattern-specific designs.

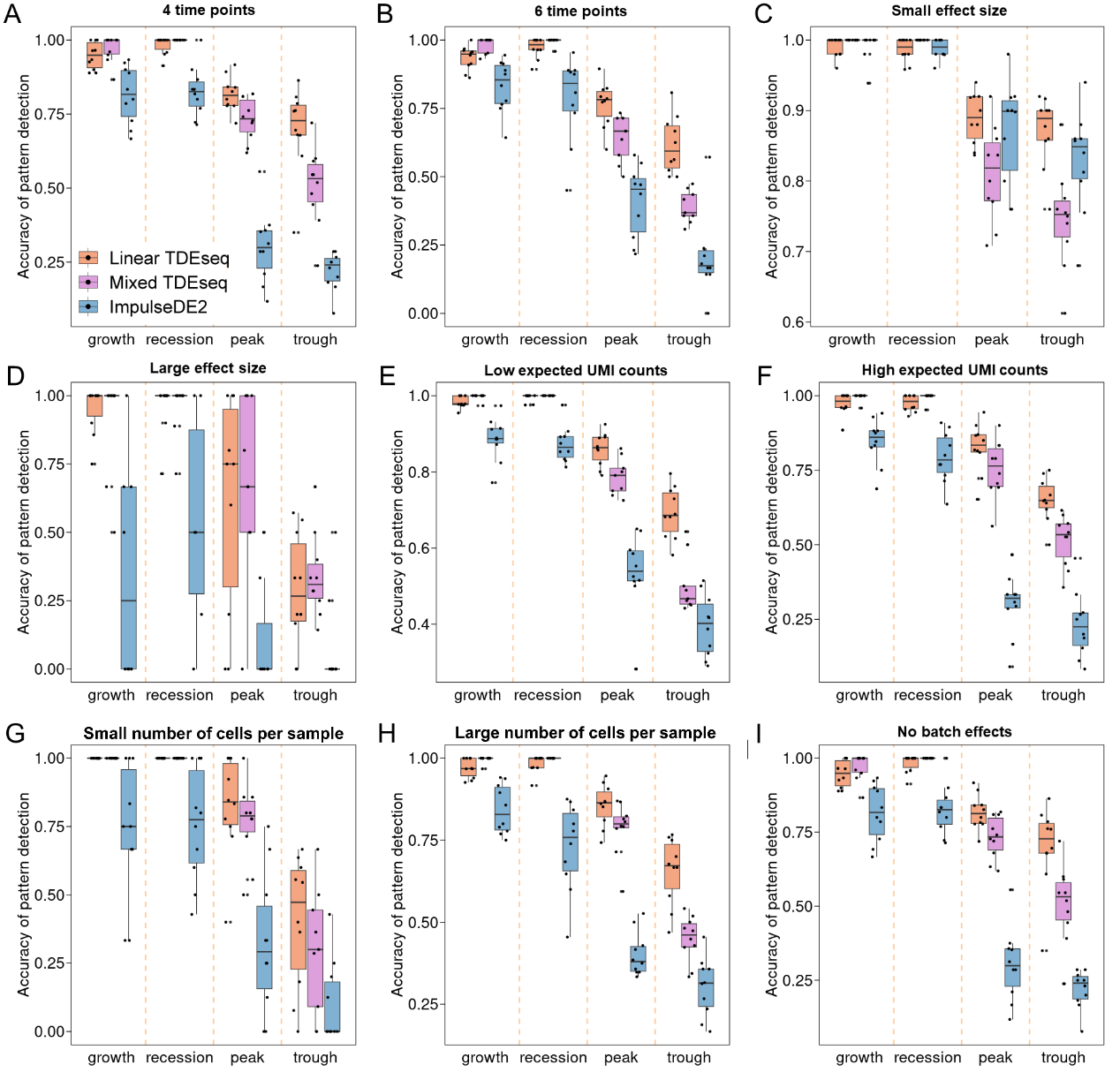

**Fig. S7.** **The comparison of the temporal expression gene detection power at an 5% of FDR in simulations.** Each simulation setup consists of 10 simulation replicates, and each replicate simulated 10,000 genes, with 1,000 temporal pattern genes and 9,000 non-temporal pattern genes. Each time point includes 3 replicates/samples. **(A)** We vary the number of time points to be either 4, 5, or 6. **(B)** We vary the effect size of biological signs to be either 0.1, 0.4, or 0.7. With increasing effect size, all methods gain power. **(C)** We vary the expected UMI counts for each cell to be either 7.0, 9.7, or 13.8. With increasing UMI counts, all methods gain power. **(D)** We vary the number of cells per individual to be either 100, 200, or 300. **(E)** We vary the batch effects to be either 0, 0.04, or 0.12. With increasing batch effects, all methods lose power. FDR: false discovery rate.

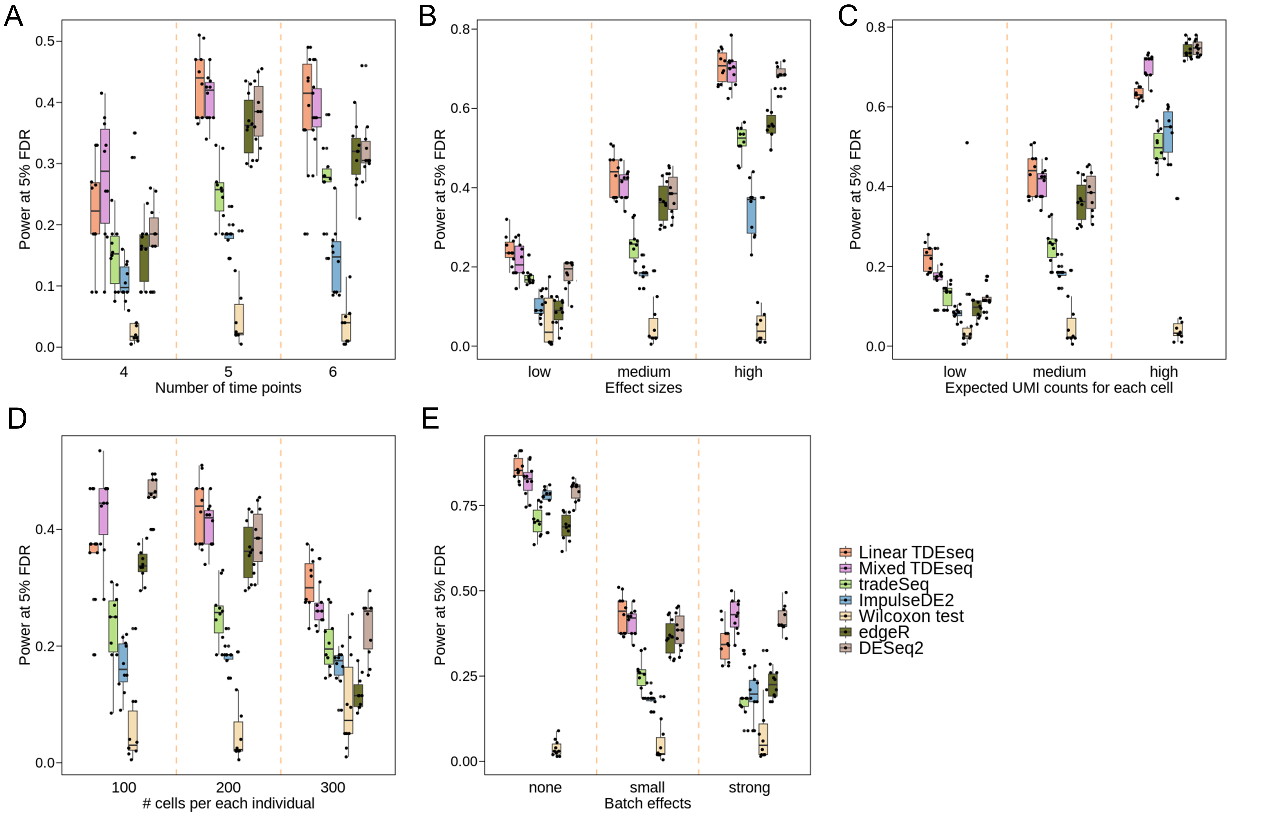

**Fig. S8.** **The performance of two additional temporal gene detection methods in simulations.** (**A**) The power comparison of temporal expression gene detection for bi-plateau pattern across a range of FDR cutoffs. (**B**) The power comparison of temporal expression gene detection for multi-modal pattern across a range of FDR cutoffs patterns. Both versions of TDEseq exhibit high detection power for detecting genes with bi-plateau expression patterns, while pseudo-bulk-based methods exhibit detection power for detecting genes with multi-modal expression patterns. The TDEseq methods were highlighted using solid lines, while other methods were represented by dashed lines in the plots. FDR: false discovery rate.

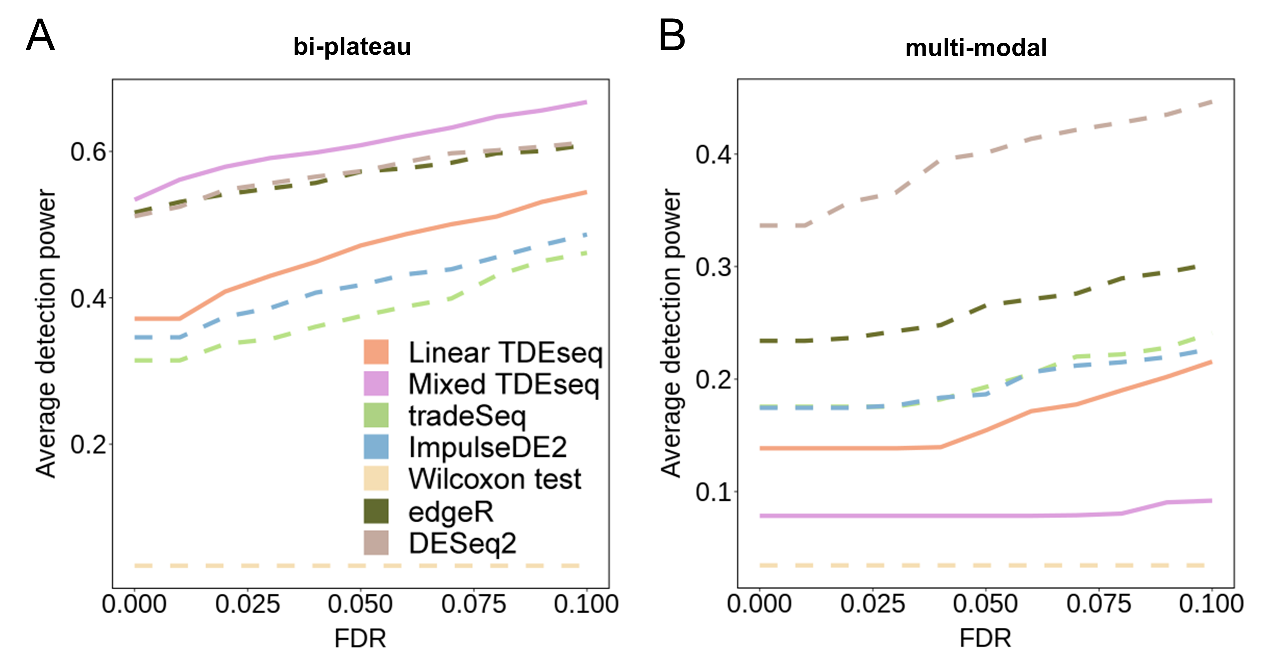

**Fig. S9. The performance of temporal gene detection methods is evaluated via varying batch effects in simulations.** (**A**) The quantile-quantile (QQ) plot shows the type I error control with no batch effects scenario. The well-calibrated *p*-values will be expected laid on the diagonal line. The *p-*values generated from Mixed TDEseq (plum), Linear TDEseq (orange), and tradeSeq (green) are reasonably well-calibrated, while ImpulseDE2 (blue), and Wilcoxon test (yellow) produced the *p*-values that are not well-calibrated. (**B**) The power comparison of temporal expression gene detection across a range of FDR cutoffs with no batch effects scenario. All temporal gene detection methods demonstrate comparable results except the Wilcoxon test. FDR: false discovery rate.

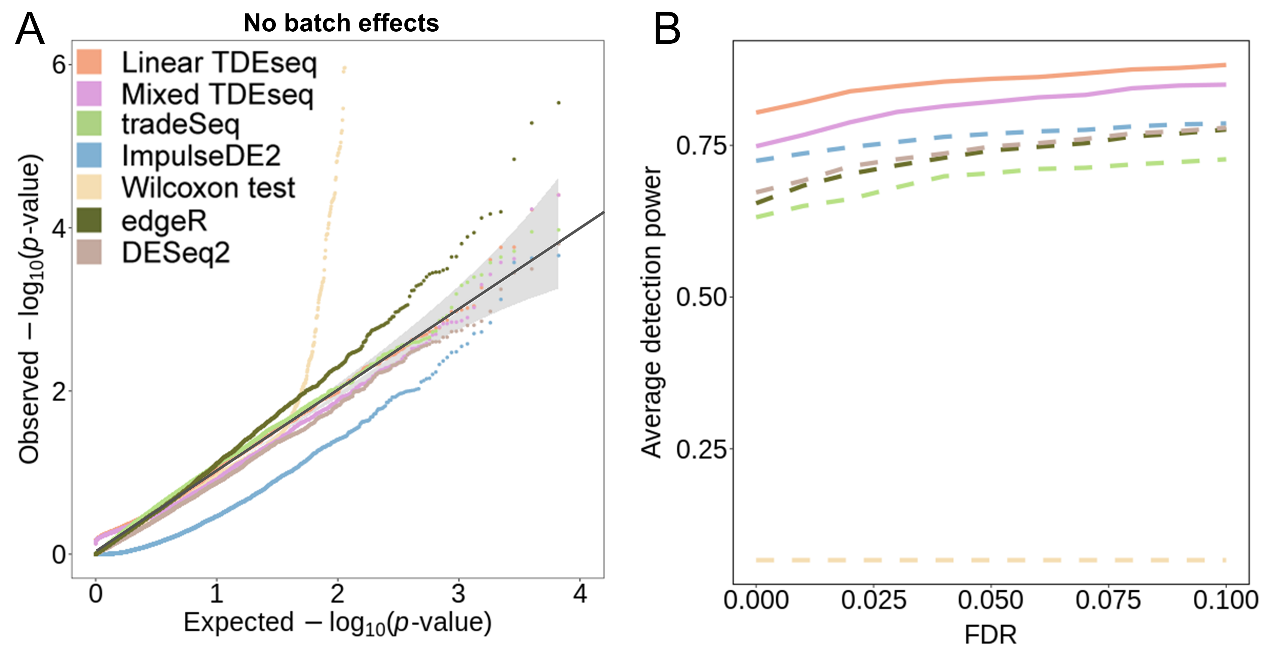

**Fig. S10.** **The UMAP visualization for the integrative analysis in simulations.** (**A**) The UMAP visualization of the original raw data. (**B**) The UMAP visualization after performing the scMerge method. (**C**) The UMAP visualization after performing MNN. (**D**) The UMAP visualization after performing Combat, (**E**) The UMAP visualization after performing Limma. (**F**) The UMAP visualization after performing ZINB-WaVE. Cells are colored by the origin of different batches. The iLISI scores were used for the evaluation of the performance of each method.

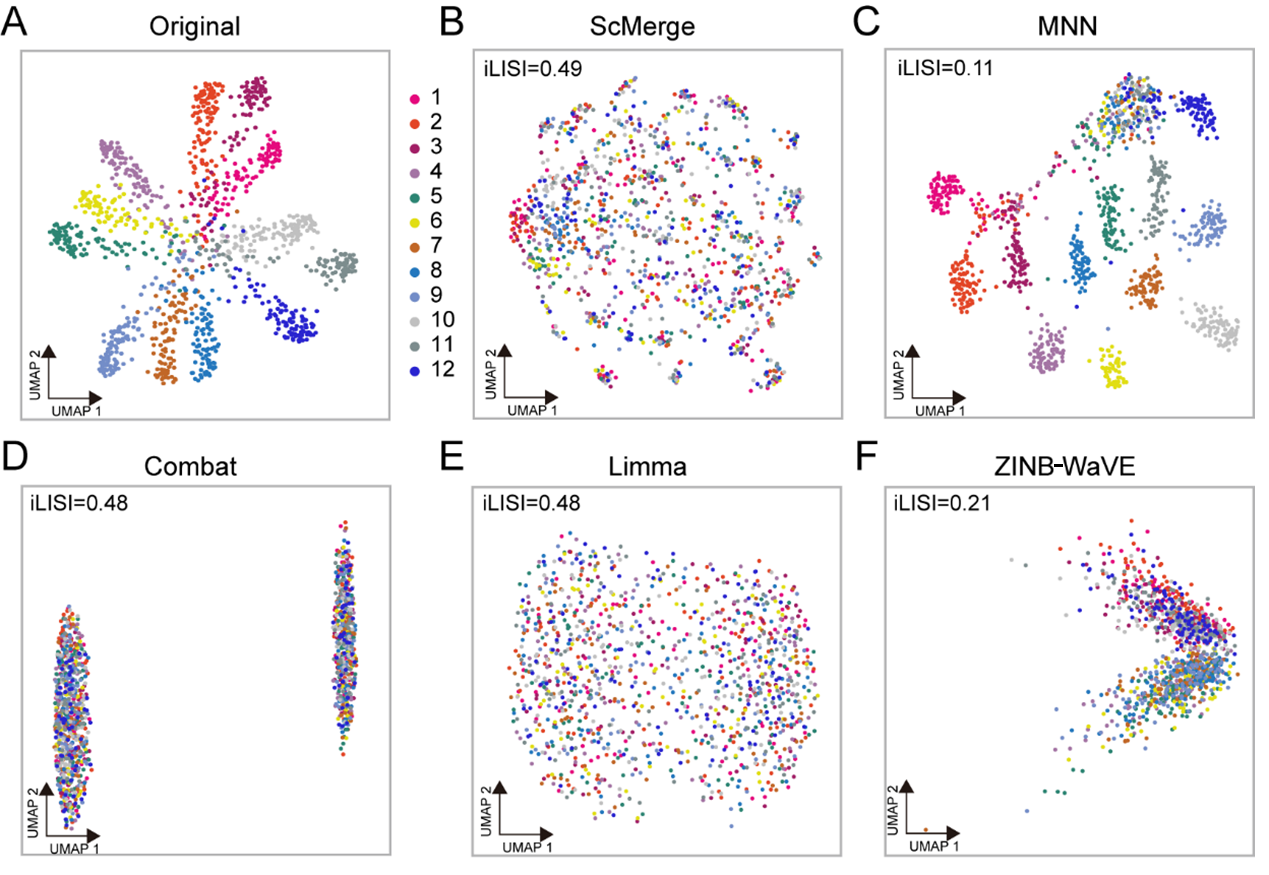

**Fig. S11.** **The performance of Mixed TDEseq coupled with different batch correction methods for large batch effects scenario.** (**A**) The type I error control is under the null. Mixed TDEseq coupled with scMerge (Mixed TDEseq + scMerge) produced approximately 6% temporal expression genes. However, Mixed TDEseq coupled with other methods produced inflated *p*-values (e.g., MNN and ZINB-WaVE) or conserved *p*-values (Limma and Combat). (**B**) The comparison of the average temporal gene detection power for Mixed TDEseq coupled with different batch correction methods, i.e. MNN (orange), scMerge (purple), ZINB-WaVE (green), Combat (blue) and Limma (azure). Mixed TDEseq + scMerge display a more powerful results of temporal expression gene detection. FDR: false discovery rate.

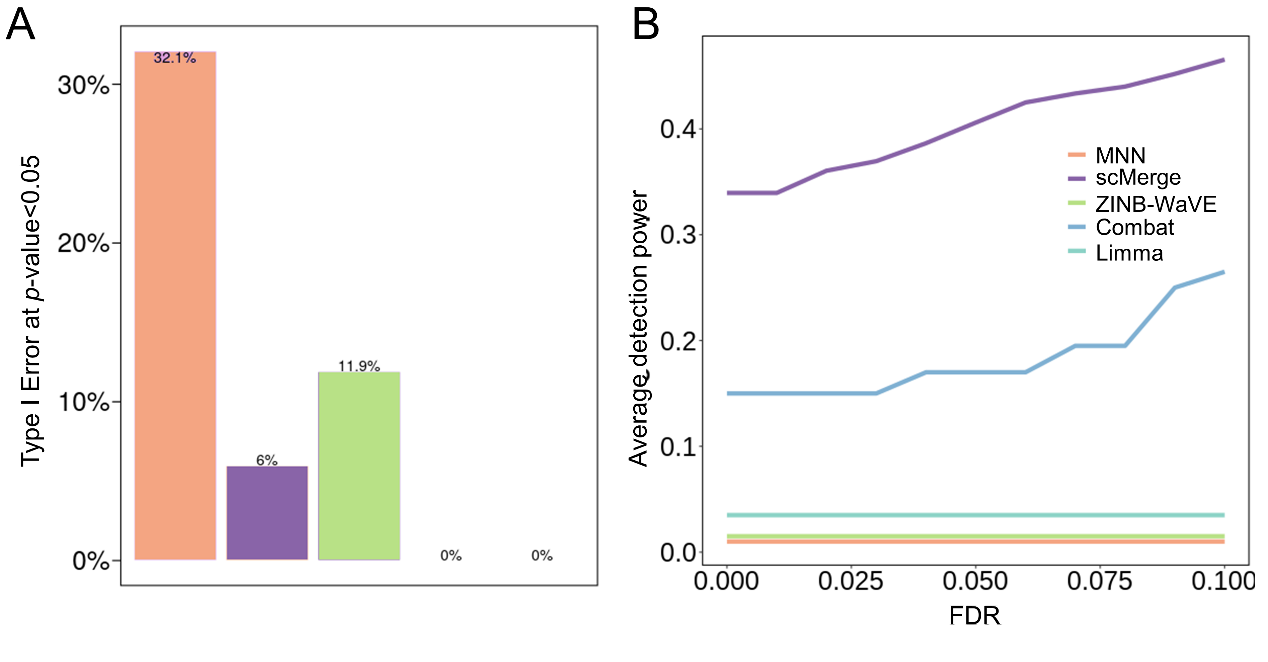

**Fig. S12.** **The performance of temporal gene detection methods is evaluated via varying the proportion of intertwined cells.** (**A**) The UMAP visualization of different proportion of intertwined cells and inferred pseudotime. The orange lines with arrows represent the direction of cell differentiation. (**B**) The power comparison of temporal expression gene detection across a range of FDR cutoffs under the small proportion of intertwined cells. (**C**) The power comparison of temporal expression gene detection across a range of FDR cutoffs under the medium proportion of intertwined cells. (**D**) The power comparison of temporal expression gene detection across a range of FDR cutoffs under the large proportion of intertwined cells. (**E**) The power comparison of temporal expression gene detection across a range of FDR cutoffs, where Linear TDEseq, Mixed TDEseq, tradeSeq, and ImpulseDE2 took inferred pseudotime as inputs. (**F**) The heatmap demonstrates the pattern-specific temporal expression genes that were identified by Linear TDEseq. Gene expression levels were log-transformed and standardized using z-scores for visualization. The temporal expression genes identified by Linear TDEseq show distinct four patterns.

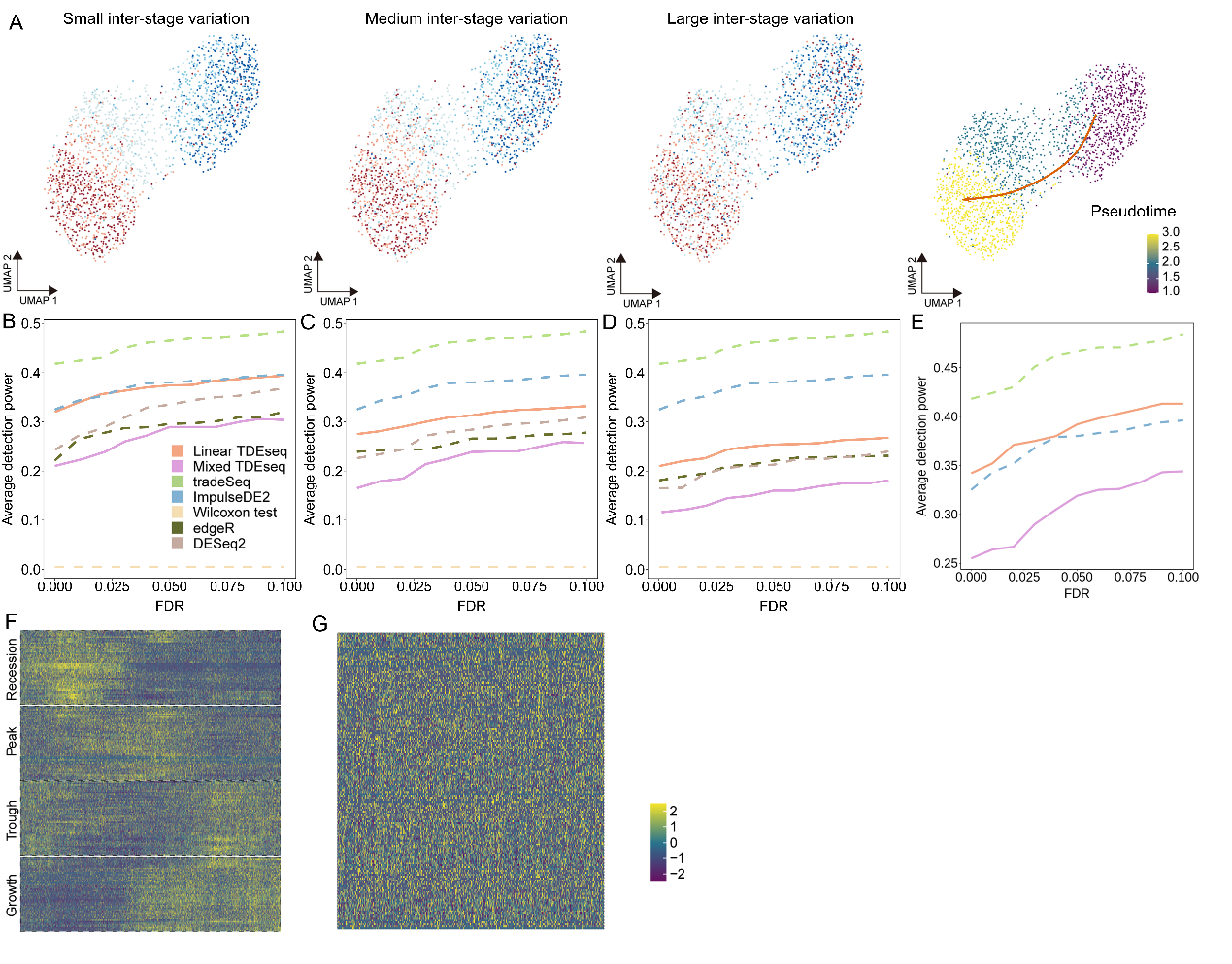

**Fig. S13. The application of scRNA-seq data from the 5-AZA-CdR treatment HCT116 cell lines**. (**A**) The UMAP visualization of low dimension of integrated 16,000 cells from 4 samples. These data sets display small batch effects over time points. (**B**) The pattern-specific detection power of Linear TDEseq (orange) and ImpulseDE2 (blue) across a range of FDR cutoffs. The TDEseq methods were highlighted using solid lines, while ImpulseDE2 was represented by dashed lines in the plots. The tradeSeq was excluded from this comparison due to no pattern-specific designs. FDR: false discovery rate.

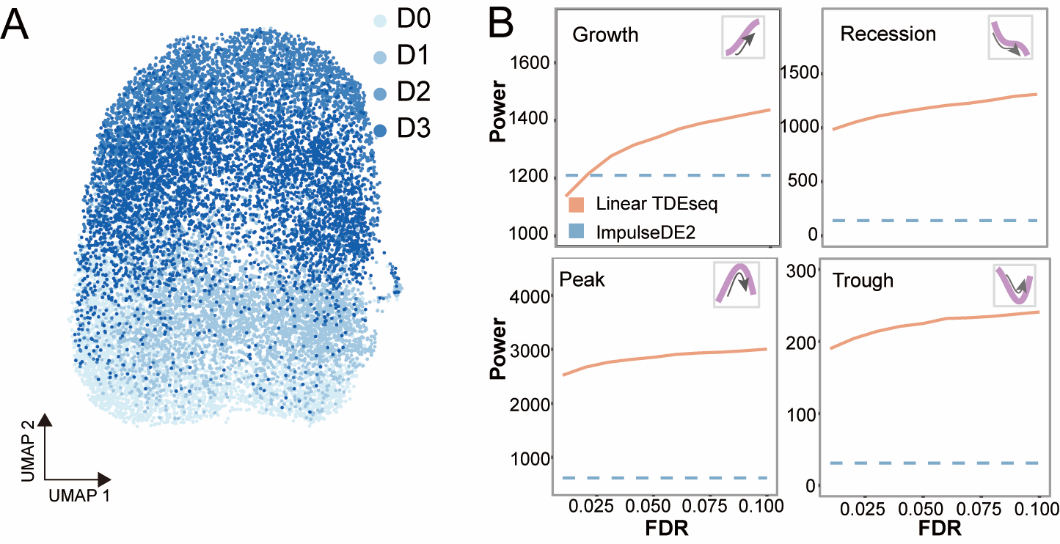

**Fig. S14**. **The application of scRNA-seq data from mouse fetal livers with seven different developmental stages**. (**A**) The UMAP visualization of integrated 13 scRNA-seq data sets from mouse fetal livers over seven developmental stages. (**B**) The pattern-specific detection power of Mixed TDEseq (plum), Linear TDEseq (orange), and ImpulseDE2 (blue) across a range of FDR cutoffs. The TDEseq methods were highlighted using solid lines, while ImpulseDE2 was represented by dashed lines in the plots. The tradeSeq was excluded from this comparison due to there being no pattern-specific designs. (**C**) The scatter density plot shows the correlation of *p*-values generated from Linear TDEseq and Mixed TDEseq. (**D**) An example gene *Mapk13* shows the difference in results between Linear TDEseq and Mixed TDEseq. Linear TDEseq only takes stage information as input and assigned a small *p*-value, while Mixed TDEseq also considered the variation among samples, and assigned a larger *p*-value. (**E**) The heatmap shows growth-specific expression genes identified by ImpulseDE2. We can see that ImpulseDE2 wrongly identified the peak or trough genes as the growth genes. (**F**) The Venn diagram shows the overlaps of temporal expression genes identified by Mixed TDEseq (plum), tradeSeq (green), and ImpulseDE2 (blue) at an FDR of 5%. The significant GO terms were enriched (*N*_GO_; BH-adjusted *p*-value < 0.05) by temporal expression genes, which were uniquely detected by Mixed TDEseq, tradeSeq, or ImpulseDE2. The temporal expression genes detected by Mixed TDEseq were enriched more GO terms. (**G**) The UMAP visualization of scRNA-seq data from mouse fetal livers with cells marked by inferred pseudotime by Slingshot. The orange lines with arrows represent the direction of cell differentiation. (**H**) The power comparison of temporal expression gene detection across a range of FDR cutoffs, where TDEseq, tradeSeq and ImpulseDE2 taken inferred pseudotime as inputs. All methods detected similar number of temporal expression genes. The TDEseq methods were highlighted using solid lines, while other methods were represented by dashed lines in the plots. (**I**) The heatmap demonstrates the pattern-specific temporal expression genes that were identified by Linear TDEseq. The top-ranked temporal expression genes identified by Linear TDEseq show distinct four patterns. Gene expression levels were log-transformed and were standardized using z-scores for visualization. FDR: false discovery rate.

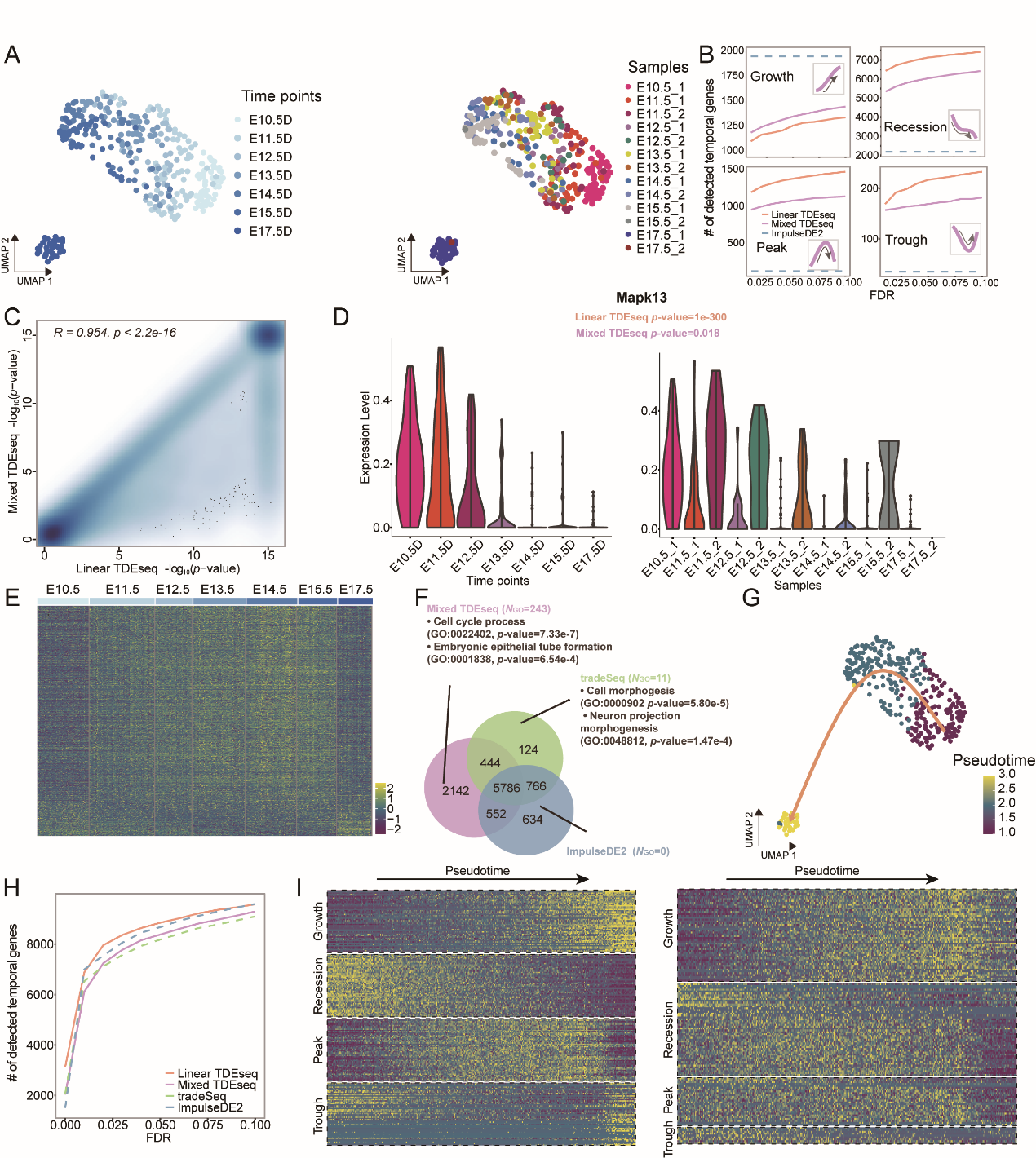

**Fig. S15**. **The application of scRNA-seq data from human metastatic LUAD with four different evaluation stages**. (**A**) The UMAP visualization of integrated 4000 epithelial cells from 26 human lung tissues over four evaluation stages. These scRNA-seq data sets display strong batch effects across different evaluation stages (left panel), and we then performed the batch effects removal using scMerge (right panel). (**B**) The pattern-specific detection power of Mixed TDEseq (plum), Linear TDEseq (orange), and ImpulseDE2 (blue) across a range of FDR cutoffs. The TDEseq methods were highlighted using solid lines, while ImpulseDE2 was represented by dashed lines in the plots. (**C**) The Venn diagram shows the overlaps of the temporal expression genes (with FDR ≤ 0.05), which were identified by Mixed TDEseq (plum), and Linear TDEseq (orange). The significant GO terms were enriched (*N*_GO_; BH-adjusted *p*-value < 0.05) by temporal expression genes, which were uniquely detected by Mixed TDEseq and Linear TDEseq. The temporal expression genes detected by Mixed TDEseq were enriched more GO terms. (**D**) The quantile-quantile (QQ) plot shows the type I error control under the permutation strategy. The well-calibrated *p*-values will be expected laid on the diagonal line. The *p-*values produced by Mixed TDEseq (plum) and Mixed TDEseq coupled with scMerge (purple) are reasonably well-calibrated. (**E**) The number of temporal expression genes identified by Mixed TDEseq (plum) and Mixed TDEseq coupled with scMerge (purple) across a range of FDR cutoffs. (**F**) The Venn diagram shows the overlaps of the temporal expression genes identified by Mixed TDEseq (plum), and Mixed TDEseq coupled with scMerge (purple) at an FDR of 5%. The significant GO terms were enriched (*N*_GO_; BH-adjusted *p*-value < 0.05) by temporal expression genes, which were uniquely detected by Mixed TDEseq or Mixed TDEseq coupled with scMerge. The temporal expression genes detected by Mixed TDEseq were enriched more GO terms. FDR: false discovery rate.

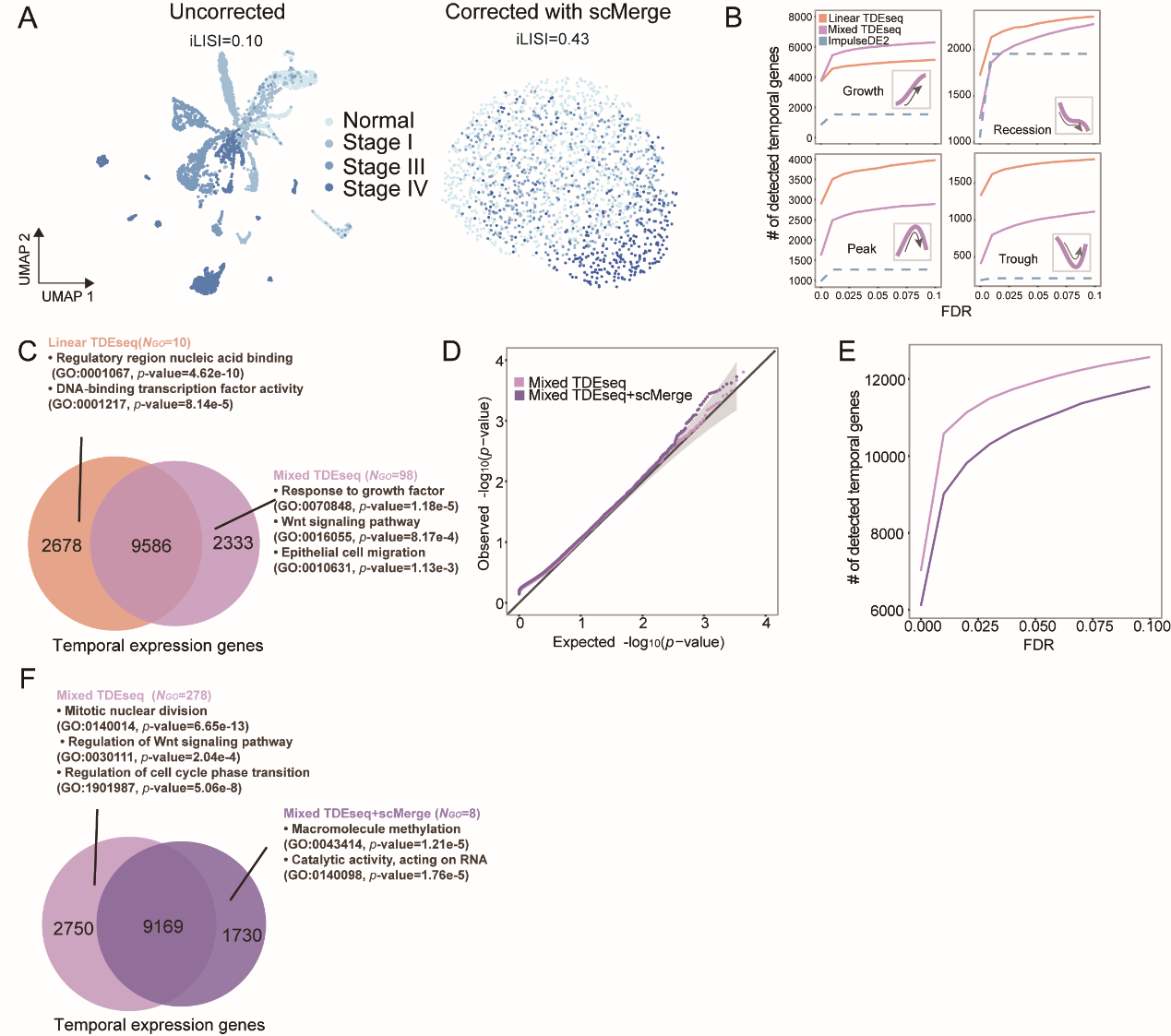

**Fig. S16**. **The application of scRNA-seq data from the NK cell response to SARS-COV-2** **infection**. (**A**) The pattern-specific detection power of Mixed TDEseq (plum) and Mixed TDEseq+scMerge (purple) across a range of FDR cutoffs. The ImpulseDE2 was excluded due to the low power. (**B**) The proportion of enrichment for the detected temporal expression genes. The given gene set (136 genes) was collected from the GO term (GO:0051607). Mixed TDEseq coupled with scMerge enriched more temporal genes than Mixed TDEseq. (**C**) The UMAP shows a growth-specific temporal expression gene, i.e., *GNLY,* which was identified by Mixed TDEseq coupled with scMerge, but missed by other methods. Another recession-specific gene*, ILF3,* was uniquely identified by Mixed TDEseq coupled with scMerge but missed by other methods*.*

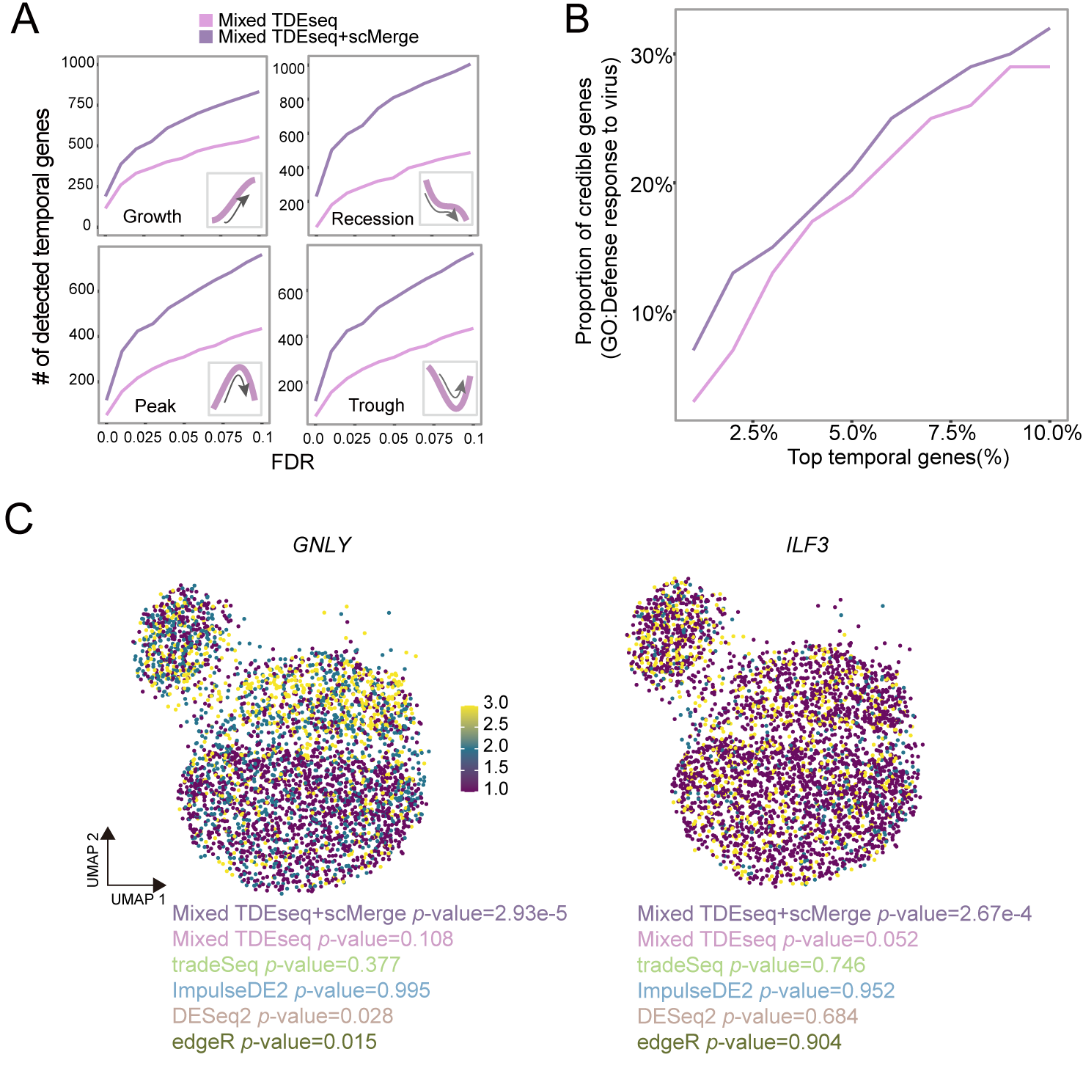

**Fig. S17. The performance of temporal expression gene detection power with varying a range of FDR cutoffs and the number of knots.** The scRNA-seq data were generated under the baseline parameter settings in simulations. The different colors denote the different number of knots. The power of TDEseq is relatively stable when the number of knots is close to the number of time points (time points = 5). FDR: false discovery rate.

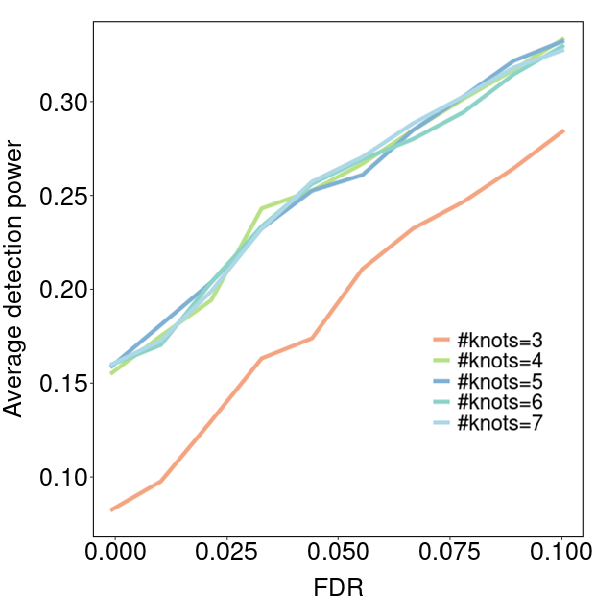
